# Epicuticular wax lipid composition of endemic European *Betula* species and its application to chemotaxonomy and paleobotany

**DOI:** 10.1101/734160

**Authors:** J. Weber, L. Schwark

**Affiliations:** Department of Organic Geochemistry, Christian-Albrechts-University, Kiel, Germany; Department of Chemistry, Curtin University, Perth, Australia

## Abstract

Plants, in particular trees with specific habitat demands are excellent indicators of climate state. Vegetation successions in subrecent and deep geologic time is recorded in fossil macro-remains or pollen accumulating in geological archives like limnic and marine sediments, peat bogs and mires. Birch trees in Europe form a major part in plant successions and constitute the dwarf species *Betula nana* and *Betula humilis* representing cold-adapted habitats or climates and two tree birches, *Betula pubescens* and *Betula pendula* characteristic for temperate habitats or climates. These birch species exhibit highly similar pollen shape and size, preventing their unambiguous application as paleoclimate/paleovegetation proxies. We here present a chemotaxonomic differentiation of the four European birch species based on their epicuticular wax lipids. The dominating lipid classes in epicuticular birch waxes were found to be n-alkanes (in the range of *n*-C_23_ to *n*-C_33_), straight-chain primary alcohols and fatty acids (in the range of *n*-C_20_ to *n*-C_32_), and long-chain wax ester (in the range of *n*-C_38_ to *n*-C_46_) in variable amounts and distributions. When preserved in geological archives these lipids may serve in paleovegetation/paleoclimate reconstruction. Long-chain wax esters are susceptible to hydrolysis and upon diagenesis the release of ester-bound alcohols and fatty acids may modify the distribution pattern of the corresponding primary free lipids. Quantitative analysis of the hydrolyzable wax ester proportion revealed primary distribution patterns of birch lipids not to change substantially upon release of bound analogues. The specific composition and abundance of epicuticular wax lipids facilitates unambiguous chemotaxonomic separation of the four European birch species. Wax lipid-based discrimination in field application, however, is complicated by mixing of alkyl lipids derived from different birch species and contribution of wax lipids from other plants. In cases, where palynology indicates a high contribution of *Betula* species to European vegetation associations, wax lipids may serve for differentiation of the species contributing.

## Introduction

Environmental demands, in particular climate, govern the present-day habitat and distribution of trees, which in turn facilitates determination of climate regimes in the contemporaneous as well as in the fossil domain. Taxonomy of present-day trees is based on anatomical, morphological, genetic and biochemical studies, whereby such features in the fossil record are best preserved in pollen distributions, due to a higher recalcitrance of pollen versus other plant organs, e.g. leaves and rare findings of other macro-remains, e.g. fruits. Under special conditions of preservation though fossil remains can be found in sediments dating back to the Eocene [1]. Paleovegetation and related paleoclimate reconstruction thus heavily relies on palynology, accompanied by analysis of marco-remains when present. Genetic [2–4] and biochemical approaches [5–8] are rare, except for molecular and isotopic investigation of leaf/needle wax, the latter rarely conducted on the species level.

Birches (*Betula* L., Betulaceae) are common broadleaf trees and shrubs occurring in diverse habitats of the boreal and the cold-temperate zones of the Northern Hemisphere [9,10]. Ranging from temperate zones in Northern America over Eurasia to East Asia and the circumpolar regions, birches populate different habitats including forests, swamps, tundra and mountainous terrains [11]. The number of species belonging to the genus *Betula* and their relationships is still under debate ranging from 30 to 60 different taxa within 4 to 6 subgenera [3,12–15]. Of these, four *Betula* species are endemic to Europe. The two tree birches, *Betula pubescens* (downy birch) and *Betula pendula* (silver birch) occur throughout most of Europe, whereby *Betula pubescens* has a more northerly and easterly distribution, while *Betula pendula* reaches more southern regions such as the Iberian Peninsula, Italy and Northern Greece [10,15]. Two dwarf/shrubby birch species, namely *Betula nana* (arctic dwarf birch) and *Betula humilis* (dwarf birch) thrive in Europe as well but are confined to a much smaller growth range. *Betula nana* is preferentially located at higher elevations like the Alps and Carpathian Mountains or in perennial colder regions like Northern Europe from Iceland over northern Scotland to Scandinavia [10,15]. *Betula humilis* has a wide distribution from western Germany to eastern Siberia and Korea, but its occurrence is very scattered with only a few habitats left in Central Europe [15]. As an important component of plant succession found in highly contrasting climate and growth regimes, these *Betulaceae* and their respective habitat demands are sensitive (paleo)climate and (paleo)environmental indicators, provided that they can be taxonomically differentiated.

Recent birch species can be distinguished by their leaf and catkin/fruit shape [16,17] even when fossilized in sediments, whereby their evolutionary relation and phylogeny is difficult to assess due to intensive hybridization in nature. Extensive hybridization and introgression has been investigated not only within a subgenus but also across *Betula* species of different subgenera [18,19]. Precise classification is complicated by an spatial overlap of natural habitats for European birch species enabling natural hybridization [20]. The identification of birch remains in geological archives for paleovegetation reconstruction is limited by a common lack of well-preserved leaves or catkins occurring in statistically relevant quantities. However, the identification of birch vegetation over time is of great interest, especially during the Late Glacial and Early Holocene (approx. 15,000 – 9,000 a BP), to understand early colonization and forest/shrub expansion in deglaciated landscapes as well as the adaptation of vegetation to climate variability and perturbation. Macrofossil findings revealed a dominance of tree birches, like *Betula pubescens*, during temperate phases, whereas upon glacial stages *Betula nana* was the most abundant birch species, serving as a tundra indicator [21].

Most commonly, paleovegetation reconstruction in sedimentary archives like lakes, peats and bogs is based on palynological approaches, since macro remains are mostly absent [22–30]. However, differentiation of birch species by pollen is challenging, due to similar morphological traits, e.g. shape, diameter and depth of pores in pollen within the genus *Betula*, which leads to overlap in pollen-size distribution [31–35]. In addition, pollen morphology can also be effected by the chemicals used upon preparation, pollen maturity, type of mounting medium, but also a latitudinal and altitudinal effect cannot be excluded (reviewed in 32). In this study we present the epicuticular leaf wax composition as a complementary proxy for recognition and differentiation of the four European birches, with the potential to employ the wax distribution patterns in paleovegetation reconstruction.

Terrestrial plant leaves are covered by a hydrophobic barrier to protect against the loss of water due to evaporation, mechanical damage, ultraviolet radiation and bacterial or fungal pathogens [36–39]. This barrier consists of an epicuticular wax layer, composed of long-chain alkyl compounds including amongst others *n*-alkanes, *n*-alcohols, *n*-alkanoic acids, *n*-alkyl esters, *n*-aldehydes, *n*-ketones and others [37,40]. Their composition is highly variable in quality and quantity across plant species and therefore has high chemotaxonomic potential [41,42,51,43–50]. *n*-Alkanes with carbon chain-length between C_25_ and C_35_ carbon atoms are associated with higher plants with a strong odd-over-even predominance (expressed as the carbon preference index-CPI), while their shorter chain homologues (<C_20_), especially *n*C_17_, are mainly found in aquatic microorganisms [52,53]. Intermediate chain-length *n*-alkanes with C_23_ and C_25_ predominance are primarily found in aquatic macrophytes and in mosses of the genus *Sphagnum* [49,54,55].

Both, *n*-alcohols and *n*-alkanoic acids in contrast to the *n*-alkanes hold an even-over-odd predominance in carbon chain-lengths, which typically ranges from C_20_ to C_32_ [37]. Alkyl esters consist of even-numbered *n*-alkanoic acids, which are esterified to even-numbered *n*-alcohols, generating long-chain aliphatic compounds with on average 38 to 52 carbon atoms [56]. Among these lipid classes, the *n*-alkanes abundances are most frequently reported in vegetation reconstruction, since they are very robust against alteration processes due to the lack of functional groups. *Betula*-derived *n*-alkanes can be found in a variety of geological archives including peats, soils, limnic and marine sediments of different ages ranging from modern times up to several million years [45,57–59] and are easily extracted from sediments and leaves by geochemical methods [37]. Therefore, most studies conducted to investigate paleovegetation history employing leaf wax lipids are based on *n*-alkanes [60]. However, the use of several lipid classes instead of a single will increase the discriminative power and representativeness of the wax lipid composition. Studies involving *n*-alkanoic acids, *n*-alcohols or *n*-alkyl ester for paleovegetation reconstruction are highly underrepresented [61,62], as these lipids are usually not reported from modern plant homologues and their degree of preservation in natural archives may vary [63]. Following incorporation into soil or sediment, wax esters can be hydrolysed, releasing free *n*-alcohols and *n*-alkanoic acids. These in turn can be converted into *n*-alkanes by decarboxylation and reduction, respectively [60]. Diagenetic fate of functionalized lipids may vary depending on various factors such as oxygen availability and pH affecting microbial reworking, but the potential of functionalized lipid classes in paleobotany has been proven for sediments of up to Miocene age [63,64].

## Material and Methods

### Leaf samples and collection

Fresh leaf samples were collected in the Botanical Garden at Kiel University in September 2017. Three leaves from each birch species were taken from branches at different sites of the tree from a height between 1 and 3 m. To avoid contaminations during sampling gloves were worn and leaves were stored in glass container or aluminium foil until extraction. Subsequently to sampling leaves were dried at 35°C in an oven for 48h. The mean annual air temperature for Kiel-Holtenau is about 9.34°C and the total annual precipitation is about 744 mm (1987-2018) (DWD, 2019)

### Lipid extraction

Lipids were extracted by immersing a single leaf sample (0.02 – 0.27 g) for 60 s in a 30-50 ml hexane/dichloromethane solution (1:1 v/v). The resulting solution was filtered through NaSO_4_, evaporated under vacuum at 50°C in a Büchi solvent evaporator and transferred into pre-weighted vials. Per species three leaves were extracted to calculate standard deviation of lipid concentration and composition. Prior to analyses, an aliquot of the total lipid extract (TLE) was treated with 35 μl *N,O*-bis(trimethylsilyl)trifluoroacetamide (BSTFA) and 5 μl pyridine at 70°C for 1 h to convert the *n*-alkanoic acids and *n*-alcohols to their corresponding trimethylsilyl (TMS) derivates. 10 μg of perdeuterated tetracosane (C_24_), octadecanol (C_18_), and eicosanoic acid (C_20_) were added as internal standard for quantification. All samples were analysed by gas chromatography-mass spectrometry (GC/MS).

### Gas chromatography-mass spectrometry (GC-MS)

The wax lipids were analysed using an Agilent 7890A (GC) equipped with an Agilent DB-5 column (30m × 0.25mm × 0.25μm) coupled to an Agilent 5975 B (MS). The oven program started at 60°C for 4 min, followed by a ramp to 140°C at 10°C/min and subsequently to 325°C at 3°C, followed by an isothermal period of 45 min. The MS operated with a scanning mass range of *m*/*z* 50-850 at an ionization energy of 70 eV. All compounds were identified by using authentic standards, NIST 14 library or their specific fragmentation pattern.

### Leaf wax characteristic calculations

The *n*-alkane content of plant species was calculated as μg/g dry weight (d.w.) of leaf based on mean values of triplicate analysis with standard deviation (Fig. 1).

**Fig 1:**
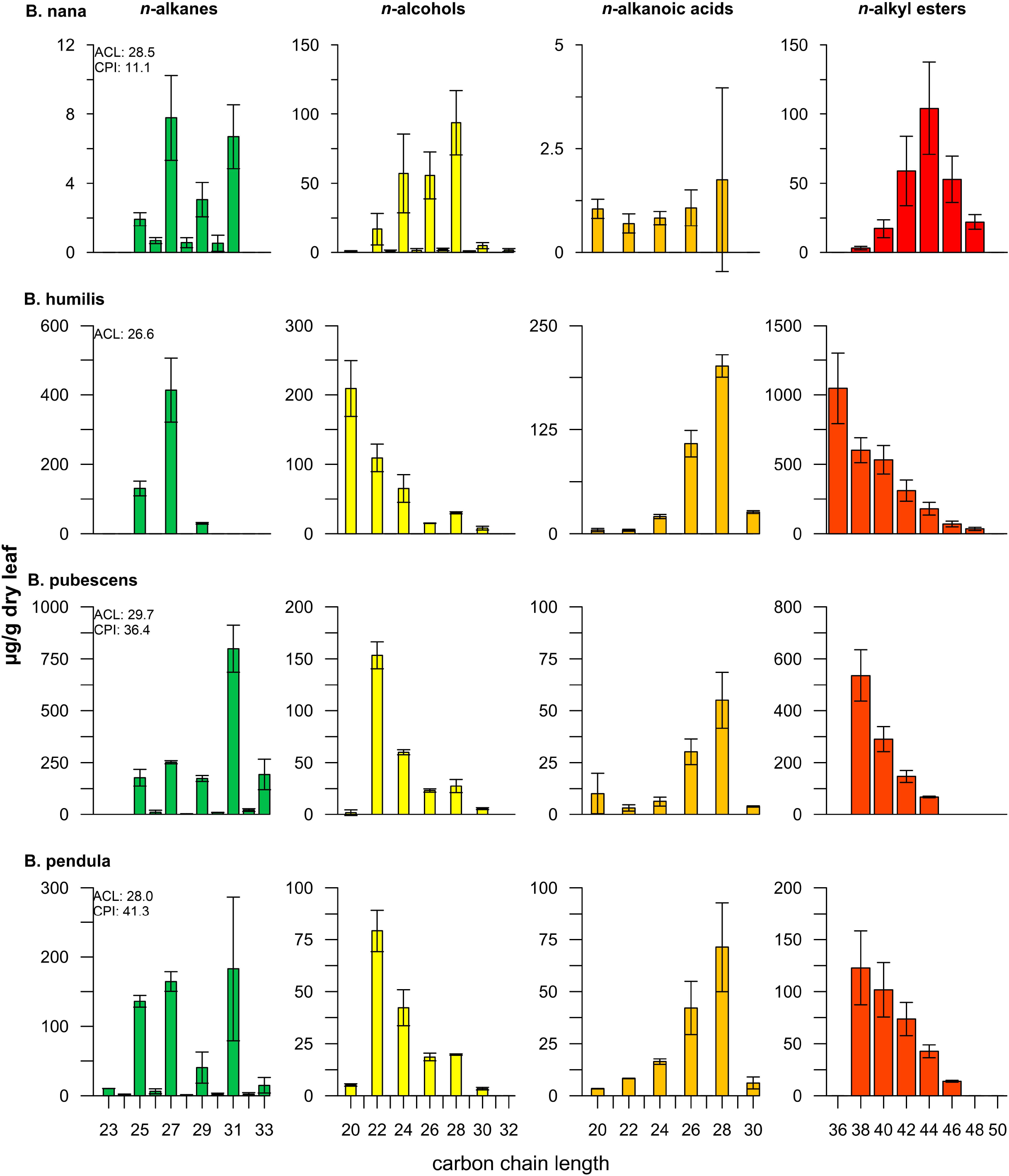
Distribution of *n*-alkanes, *n*-alcohols, *n*-alkanoic acids and *n*-alkyl esters (μg/g dry leaf) from epicuticular waxes of four European birch species collected the Botanical Garden of Kiel (Germany). Note the different scale of y-axis for each row.

Average chain length (ACL) for *n*-alkanes with 23 to 33 carbon atoms was calculated as:

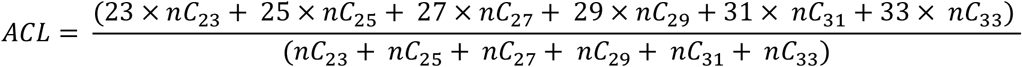

with *C_n_* as relative abundance of *n*-alkanes with the chain length *n* [65]. This proxy is used as weighted mean of *n*-alkane carbon chain length, which supposed to vary with climate. The carbon preference index (CPI) outlines the relative abundance of odd-over-even carbon chain lengths, whereby values >1 indicate a predominance of odd carbon chain lengths homologues. CPI values for *n*-alkanes with 24 to 34 carbon atoms were calculated according to:

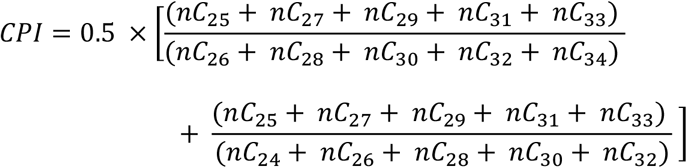

### Wax ester quantification

For total wax ester quantification, the respective wax ester peaks in the GC-MS chromatograms were integrated and quantified against an internal standard (deuterated tetracosane). For a wax ester of the type RCOOR’ the diagnostic fatty acid ion is RCOOH_2_^+^, indicative for the alkanoic acid chain length. R’ – 1^+^ derives from the corresponding alcohol moiety, however, its intensity is low and difficult to detect. Therefore, the diagnostic acid fragments (RCOOH_2_^+^) of peaks containing co-eluting alkyl esters of identical total mass but variable combinations of alcohol and alkanoic acid moieties were integrated to determine the percentage of the respective isomer contribution. Multiplication of isomer percentages by analogue abundances led to the proportion of the esterified acids. The proportional amount of esterified alcohol was obtained by subtracting the esterified acid from the corresponding total wax ester homologue.

## Results and discussion

### Alkyl lipid distribution of plants from Botanical Garden of Kiel University

The focus of this study lies on the epicuticular wax composition of the four birches endemic to Europe. The leaf wax compound classes *n*-alkanes, *n*-alcohols, *n*-alkanoic acid and *n*-alkyl ester were present in all species at variable composition distributions (S1). Epicuticular alkyl lipid abundances are reported as μg/g dry weight (d.w.) of leaf.

The lowest total amount of epicuticular waxes was observed in the artic dwarf birch *B. nana* with 538.3 μg/g d.w., followed by *B. pendula* with 1293.8 μg/g d.w., 3131.7 μg/g d.w. in *B. pubescens* and *B. humilis* with 4187.9 μg/g d.w..

### *n*-alkane of *Betula* epicuticular wax

The carbon atom chain-lengths of *n*-alkanes varied from *n*C_23_ to *n*C_33_ and maximized either at *n*C_27_ or *n*C_31_. A predominance of nC_27_ was noted for *B. humilis* with 413.2±91.1 μg/g d.w.. In relative proportion, nC_27_ was the dominant *n*-alkane homologue in *B. nana*, though it occurred in minor absolute concentrations (7.8±2.5 μg/g d.w.) only. The *n*-alkanes of two tree birches, *B. pubescens* and *B. pendula*, maximised at nC_31_ with 799.0±133.3 and 183.0±103.8 μg/g d.w., respectively, whereby the latter species additionally contained large proportions of *n*C_25_ and *n*C_27_ alkanes. The average chain-lengths for odd-carbon-numbered *n*-alkanes in the range from *n*C_23_ to *n*C_33_ (ACL_23-33_) varied from 26.7 in *B. humilis* to 29.7in *B. pubescens*, while *B. pendula* und *B. nana* had intermediate ACL values of 28.0 and 28.5, respectively. As expected, in the recent leaves a typical odd-over-even predominance was detected, which is expressed in high CPI values. Highest CPI values were found in *B. pendul*a averaging 41.3, followed by *B. pubescens* with 36.4 and *B. nana* with 11.1. The CPI for *B. humilis* could not be calculated due to the lack of even-numbered *n*-alkanes. The *n*-alkane CPI values of the three species were high, indicating that there was no significant contamination by diagenetic or petroleum-derived *n*-alkanes.

The birch trees examined in this study grew under identical environmental conditions in the Botanical Gardens of Kiel University, thus a possible influence of climate or soil condition on the wax lipid distribution should be identical for each species. The *n*-alkane compositions in all four Betula species exhibited a dominance of longer-chain lengths homologues (*n*C_27_, *n*C_29_, *n*C_31_) with a mean ACL of ca. 28, as typical of deciduous trees [49]. However, some species had only one dominant homologue (*B. pubescens, B. humilis*), whilst others had a more bimodal distribution with two dominant homologues (*B. nana, B. pendula*) (Fig. 1). The ACL has been used in geological archives as a climate and plant species indicator. For example, it has been recognized that higher ACL values correlated with higher temperatures in sediments of Lake Malawi (east Africa), potentially as protection against heat to increase the melting point of epicuticular waxes [66]. However, Diefendorf et al. 2017 in their compilation study could not confirm a significant temperature control on ACL neither in C_3_ woody plants nor in C_3_/C_4_ grasses. Higher ACL values, due to a higher proportion of *n*C_31_, *n*C_33_ and partially *n*C_35_, were observed in C_4_ grasses (*poaceae*) of arid zones in Africa, which distinguished them from C_3_ species from Peru and Australia [67]. A correlation between the preferred habitat of the four birches from our study (arctic/alpine *vs*. temperate zone) and ACL is not noticeable, since especially the two cold tolerant dwarf birches showed markedly different *n*-alkane distributions. However, the measured ACL values were in accordance with the expected values, since no extended long-chain *n*-alkanes with 33 or even 35 carbon atoms, typically for arid grasses, were found.

### *n*-alcohols of *Betula* epicuticular wax

As with the *n*-alkane distribution, a typical terrestrial higher plant pattern was observed, yielding a strong even-over-odd dominance in carbon chain-lengths [68]. The *n*-alcohol chain-lengths ranged from *n*C_16_ to *n*C_32_ with a predominance in long-chain alcohols (>*n*C_20_) in all birches. As short chain-lengths *n*-alcohol homologues (<*n*C_20_) are primarily synthesised by microbes or algae [69], we based epicuticular wax analysis on the long-chain homologues. The primary *n*-alcohols showed the closest range in overall concentrations of all lipid classes (Fig. 1). The cumulative *n*-alcohol concentration extended from 168.5±22.0 μg/g in *B. pendula* to 436.4±84.7 μg/g in *B. humilis*. The three birches *B. humilis, B. pubescens* and *B. pendula* revealed a gradually decreasing concentration with increasing chain-lengths, whereby *B. humilis* maximized at *n*C_20_ with 209.0±40.2 μg/g d.w. and the latter two at *n*C_22_ with 153.5±12.9 and 79.2±10.0 μg/g d.w., respectively. *B. nana* was characterized by a narrower distribution, peaking at *n*C_28_ with 99.7±23.3 μg/g d.w.. Our results demonstrate that the concentration of *n*-alcohols either increased or decreased gradually with increasing carbon chain-length with only one homologue being dominant (Fig. 1). Both tree birches, *B. pubescens* and *B. pendula*, revealed similar alcohol patterns and differed in absolute concentrations only.

In contrast to the tree birches, however, the two shrub birches showed a markedly different *n*-alcohol distribution and were well distinguishable.

Diefendorf et al. (2011, 2015) reported that the average concentrations of *n*-alcohols in leaves of deciduous angiosperm trees from the U.S. commonly were twice as high as those of *n*-alkanes. In our study *B. nana* exclusively revealed about 10 times higher concentrations of *n*-alcohols compared to *n*-alkanes. The other three species produced two to four times lower amounts of *n*-alcohols than *n*-alkanes. Previous studies have shown that grasses are mainly characterized by *n*C_26_, *n*C_28_ and *n*C_32_ alcohols [67,70]. This corresponds closely to generally low amounts of these homologues in favour of the high proportions of the shorter alcohols *n*C_20_ or *n*C_22_ in *B. humilis*, *B. pubescens* and *B. pendula*. Only the leaves of *B. nana* produced a higher proportion of *n*C_26_ and *n*C_28_ alcohols.

### *n*-alkanoic acids of *Betula* epicuticular wax

The *n*-alkanoic acid abundances were characterized by a strong even-over-odd dominance in carbon chain-lengths in the range of *n*C_12_ to *n*C_30_. Similar to the *n*-alcohols, short-chain homologues (<*n*C_20_) were not significant for higher plants since these compounds are also produced by a variety of organisms like bacteria and algae, or derive from cellular membranes rather than waxes. Here, in all four birches species alkanoic acids in the range of *n*C_12_ to *n*C_30_ were observed to peak at *n*C_16_ or *n*C_28_. In the range of the long-chain fatty acids (>*n*C_20_), the four Betula species maximized exclusively at *n*C_28_, whereby their concentrations differed by two orders of magnitude (Fig. 1). Thus, *B. nana* yielded only 1.75±2.2 μg/g d.w. of *n*C_28_, while *B. humilis* produced 201.2±13.4 μg/g d.w. of C_28_. *B. pubescens* and *B. pendula* showed intermediate concentrations of 34.3±2.4 and 34.7±4.0 μg/g d.w., respectively. *B. nana* produced more short chain alkanoic acids than long-chain homologues, with highest concentration at *n*C_16_ with 44.3±6.1 μg/g d.w. and *n*C_18_ with 20.8±1.3 μg/g d.w..

The relative distribution patterns of long-chain alkanoic acids in the four birches were too similar to allow for differentiation. These basic findings were consistent with a litter and topsoil transect experiment in which deciduous forest sites also showed a dominance of *n*C_28_ alkanoic acids and differed from conifer (*n*C_24_) and grasslands sites (*n*C_32_ and *n*C_34_) [71]. Therefore, C_28_ alkanoic acid preponderance may serve to distinguish wax lipid inputs of birches from those of grasses and conifers, like pines. This may be applicable for sediments in periods such as the Late Glacial and Early Holocene in Central Europe, where successions were characterised by a minor diversity in plant species. In contrast, Diefendorf et al. (2011) observed that the *n*-alkanoic acid distributions across plant groups from the east coast of the USA, both angiosperms and gymnosperms, were similar with no dominant homologue present.

### *n*-alkyl esters of *Betula* epicuticular wax

Wax esters are dimeric wax compounds build by a *n*-alcohol and a *n*-alkanoic acid moiety, whereby each *n*-alkyl ester isomer can be composed of several different combinations of *n*-alcohol and *n*-alkanoic acid homologues (S2) [72]. Saturated wax ester homologues were identified according to their characteristic molecular ions (M^+^) [73]. Straight-chain wax esters in the range of *n*C_38_ to *n*C_48_ were detected in all four species, which is typical for higher plants [74] (Fig. 1). Additionally, *B. humilis* produced minor quantities of the *n*C_36_ homologue. The alkyl ester composition of *B. nana* maximized at *n*C_44_ with 104.2±33.4 μg/g d.w. with an almost normal distribution. *B. humilis* peaked at *n*C_36_ with 1047.1±254.0 μg/g d.w., followed by a linear decrease in concentration of longer-chain wax esters up to *n*C_48_. Wax esters chain lengths of *B. pubescens* and *B. pendula* leaves ranged from *n*C_38_ to *n*C_46_, whereby the concentrations decreased with an increase in chain-lengths. Concentrations of *n*C_38_ wax ester of *B. pubescens* and B. pendula yielded 535.7±99.4 and 122.7±35.5 μg/g d.w., respectively.

### Isomer distribution of wax esters and input to free *n*-alkanoic acids and *n*-alcohol amount

The mass spectral analysis of wax esters by GC/MS allowed to investigate their corresponding bound *n*-alcohol and *n*-alkanoic acids (Fig. 6, S3–S6 Tables).

In all four species, only even-chain alkanoic acids in the range of *n*C_14_ to *n*C_28_ and alcohol moieties ranging from *n*C_18_ to *n*C_32_ were observed resulting in even-chain alkyl esters. In *B. nana*, *n*C_20_ was the overall dominant ester-bound alkanoic acid moiety with 62.2 μg/g d.w. and *n*C_24_ the most prominent ester-bound alcohol with 58.9 μg/g d.w.. In the shorter *n*C_38_ and *n*C_40_ esters shorter ester-bound alkanoic acids with 14 and 16 carbon atoms as well as shorter ester-bound alcohols like *n*C_22_ occurred. The ester homologues of *B. humilis*, *B. pubescens* and *B. pendula* were dominated by short-chain *n*C_16_ bound alkanoic acid moieties (841.5, 368.8, 75.7 μg/g d.w.), while *n*C_20_ was the most dominant alkanoic acid homologue in the long-chain alkanoic acid fraction (190.6, 38.0, 33.4 μg/g d.w.). However, compared to the short-chain alkanoic acids, the long-chain homologues were less abundant in concentrations up to factor of 10. The major esterified alcohol within the three species varied significantly. *B. humilis* was dominated by *n*C_20_ (744.2 μg/g d.w.), *B. pubescens* by *n*C_22_ (346.6 μg/g d.w.) and *B. pendula* by *n*C_24_ (82.7 μg/g d.w.) and *n*C_22_ (81.2 μg/g d.w.) bound alcohols.

The bound *n*-alkanoic acid and *n*-alcohol moieties of wax esters might be released during hydrolysis upon incorporation of alkyl esters into soil or during early burial stages in sediments. As consequence, the amount of hydrolysis-released, previously ester-bound *n*-alkanoic acids and *n*-alcohols in a sediment will impact on the quantity and distribution pattern of free *n*-alkanoic acids and *n*-alcohols derived from leaf waxes [70] and needs to be considered in paleovegetation reconstruction. However, intact wax esters can survive in sediments and can be used for paleovegetation reconstruction [63,75,76] as well.

To test for the potential release of alcohols and acids from esters two different scenarios were calculated. In the first scenario, 50% of the esterified *n*-alkanoic acids and *n*-alcohols were released and added to their corresponding free homologues (Fig. 2). In the second scenario, the maximum release of 100% of the bound lipids and addition to the free analogues was used. Since the wax esters of all four birches consisted mainly of short-chain fatty acids, they do not significantly increase the pool of long-chain fatty acids (>*n*C_20_) typical for terrestrial higher plants. Solely in *B. nana*, the alkanoic acid distribution changed from a previously rather balanced distribution of *n*C_20_ to *n*C_28_ with 1 to 2 μg/g d.w. to a dominance of *n*C_20_ with >60 μg/g d.w., due to the release of the esterified homologues. A dominance of *n*C_28_ alkanoic acid was still observed for the other three birches when 50% of the esterified alkanoic acids had been released. Only upon 100% release of the bound long-chain fatty acids, a bimodal distribution maximizing at *n*C_20_ and *n*C_28_ could be noted for *B. pubescens*, which would complicate a source identification in sediments.

**Fig. 2:**
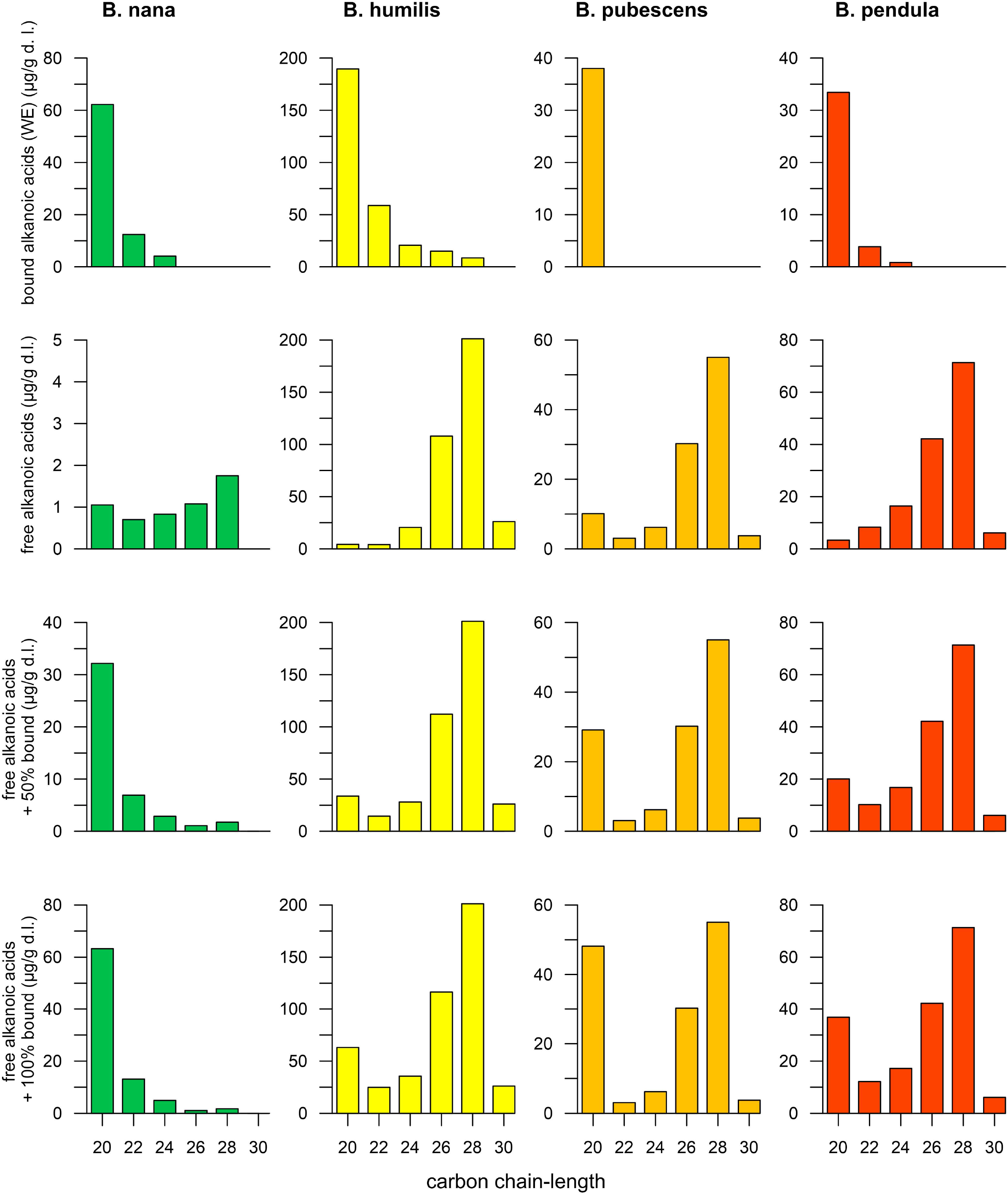
Distribution of esterified (bound) alkanoic acids in the alkyl esters and their corresponding free homologues in the same leaf. The two lower rows depict the summed concentration of free fatty acids plus an additional 50% or 100 % bound fatty acids released by hydrolysis, respectively.

The bound *n*-alcohols of the alkyl esters in the four European birches ranged from *n*C_20_ to *n*C_32_. Sometimes, the dominant esterified *n*-alcohol was found to be the same as the dominant free homologue in the same species (Fig. 3). For example, in *B. humilis n*C_20_ and in *B. pubescens n*C_22_ were the dominant free and esterified *n*-alcohol, respectively. The release of 100 % bound *n*-alcohols of both species increased the total amount by about 25%. However, the relative distributions remained comparable. The previously identified dominance of the free *n*C_22_ alcohol in *B. pendula* was reduced by the release of a high proportion of *n*C_24_. In contrast, the distribution in *B. nana* changed from a dominance of *n*C_28_ to a bimodal distribution maximizing at *n*C_24_ and *n*C_28_ due to the addition of ester-bound *n*-alcohols. The dominance of free *n*C_22_ alcohol in *B. pendula* was reduced by the release of the high proportion of *n*C_24_.

**Fig. 3:**
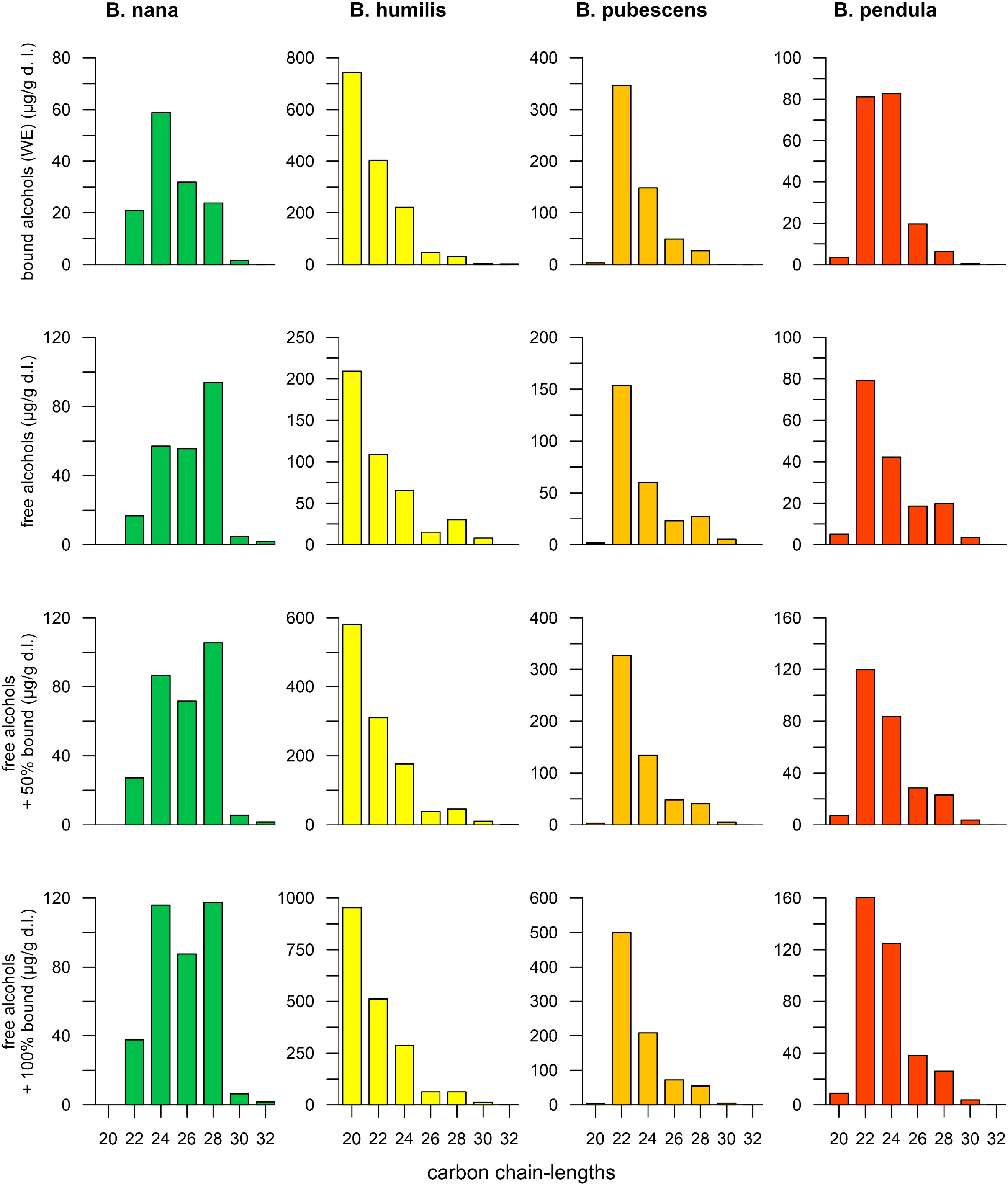
Distribution of esterified (bound) alcohols in the alkyl esters and their corresponding free homologues in the same leaf. The two lower rows depict the summed concentration of free fatty acids plus an additional 50% or 100 % bound fatty acids released by hydrolysis, respectively.

Our model indicates that the decay of the *n*-alkyl ester can significantly affect the original free lipid composition of birch leaves, which in sediments may complicate an unambiguous assignment based on *n*-alcohol and *n*-alkanoic acids.

### Wax lipids from *Betula* grown in Kiel compared with literature data

To consolidate the application of the lipid composition of the four European birches for paleovegetation reconstruction and chemotaxonomic differentiation, we compared our wax lipid data of birch trees grown in the Botanical Garden in Kiel with previously published data (Table 1). However, solely the *n*-alkane composition of *B. nana, B. pubescens* and *B. pendula* can be compared, as to the best of our knowledge, the other wax lipid classes were not examined in full in previous studies. To our best knowledge, there are no previous studies on the wax lipid composition of *B. humilis* leaves, thus we do not have any complementary data for comparison. For a better comparison of published data, we have recalculated the distributions given in absolute concentrations into relative abundances, to exclude variation in absolute concentration due to different extraction techniques or analytical protocols. Different extraction techniques and analytic procedures used in the cited studies are briefly described here:

- Extraction of epicuticular waxes by immersion of the hole leaf into solvent mixture with or without ultrasonic bath, e.g. DCM:hexane (1:1) or pure DCM (our study, [77]);
- Prior to extraction, grounding or milling of leaves into a fine powder, followed by ASE (extraction under elevated temperature and pressure) or Soxhlet extraction [78–80]; here, intra-cuticular waxes potentially have been extracted;
- Hydrolysis (10% KOH in ethanol) of ground leaves to extract bound lipids [81]; esterified alkanoic acids and alcohols were released and increased the pool of their free homologues.

**Table 1:**
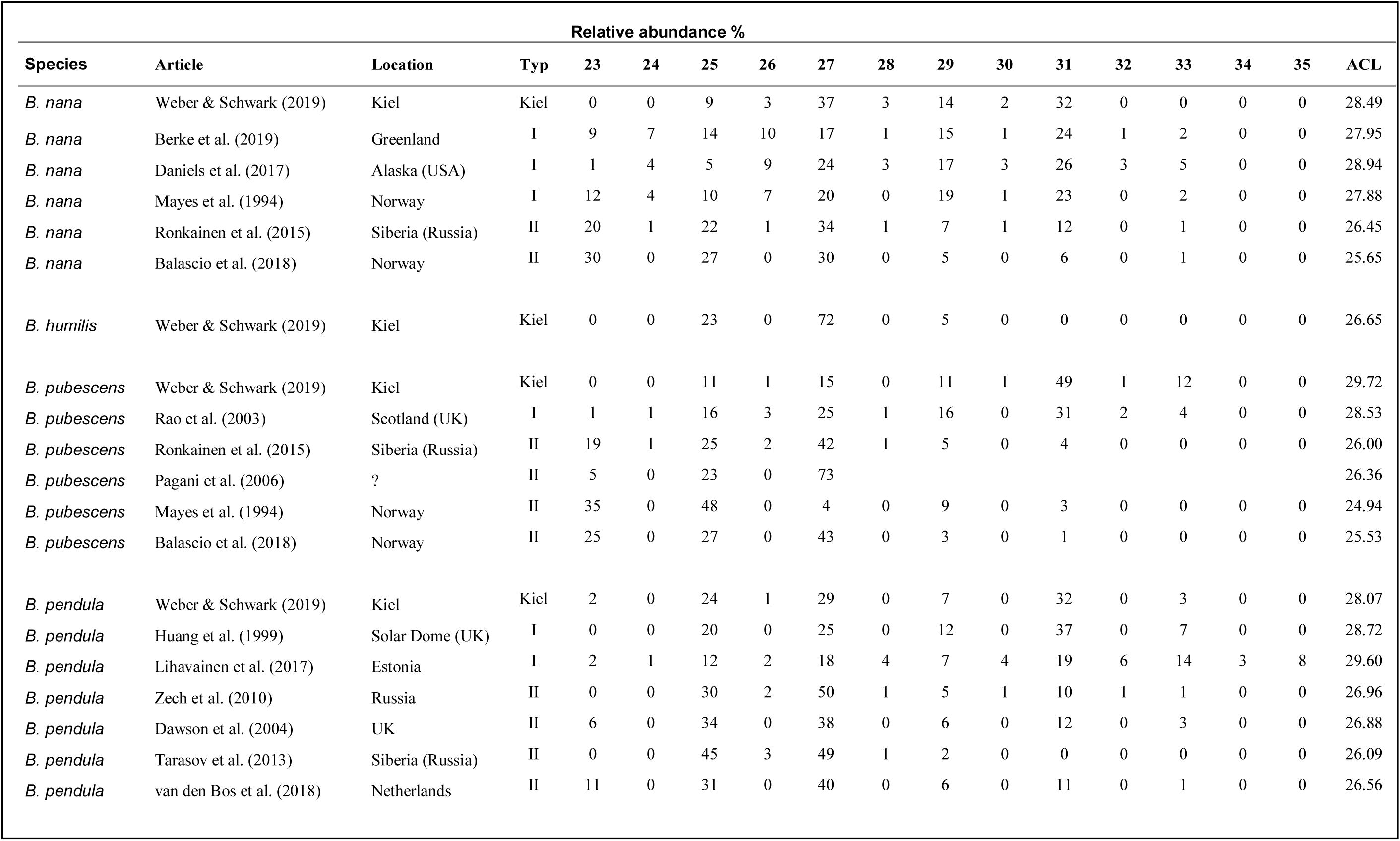
Relative *n*-alkane abundances of all *Betula* species

Most leaves from B. *nana* are characterized by a variable distribution of *n*-alkanes ranging from *n*C_23_ to *n*C_33_. Samples from very northern latitudes such as Greenland, Alaska, Siberia and Norway had a high proportion of *n*C_27_ to *n*C_31_ homologues [80–82]. Conversely, *B. nana* leaves from Siberia and Norway can be distinguished from the other *B. nana* samples as these showed significant abundances of mid-chain *n*C_23_ and *n*C_25_ alkanes depressing relative amounts of C_29_ and C_31_ homologues [83,84]. Leaf *n*-alkanes of *B. pubescens* from Scotland were bimodally distributed with maxima at *n*C_27_ and *n*C_31_ [85]. In contrast, a unimodal distribution with a maximum at *n*C_27_ was reported from Pagani et al. (2006), Ronkainen et al. (2015), and Balascio et al. (2018), with variable proportions of *n*C_23_ and *n*C_25_, respectively. A dominance of C_27_ alkane in leaves from *B. pubescens* has also been reported from Schwark et al. (2002) and Sachse et al. (2006). The latter author stated that only in birches (both *B. pubescens* and *B. pendula*) from northern Scandinavia *n*C_27_ is the dominant homologue, while species from southern Scandinavia and Germany maximized at *n*C_31_. This shift in *n*-alkane carbon chain-length was also expressed in an increasing ACL from North to South [87]. In contrast to this, Mayes et al. (1994) found a prevalence of C_25_ in leaves from Norway.

Similar to the leaves of *B. pubescens*, those of *B. pendula* contained high proportions of C_25_, if the distribution maximized at *n*C_27_ and then constituted only minor long-chain homologues with more than 29 carbon atoms [78,79,88,89]. Two *B. pendula* species from Estonia and the UK, when grown under artificial laboratory conditions had a distinct bimodal distribution with maxima at C_27_ and C_31_ [77,90].

Due to the complexity of the *n*-alkane patterns both within a species and between different species, we subdivided the data according to distributions of *n*-alkanes in each species into three groups to improve comparison. The published data of each species were subdivided into two groups (type I and type II) of similar composition and compared with the lipid distribution of the birches from Kiel (Fig. 4, Table 2). Type I of each species had a wax lipid composition similar to the *Betula* species from Kiel and was characterized by a dominance of long-chain *n*-alkanes (*n*C_27_, *n*C_29_, *n*C_31_). In contrast, type II of each species was defined by a high presence of mid-chain *n*-alkanes (*n*C_23_, *n*C_25_), but also *n*C_27_, and only minor quantities of long-chain *n*-alkanes with more than 29 carbon atoms.

**Figure 4:**
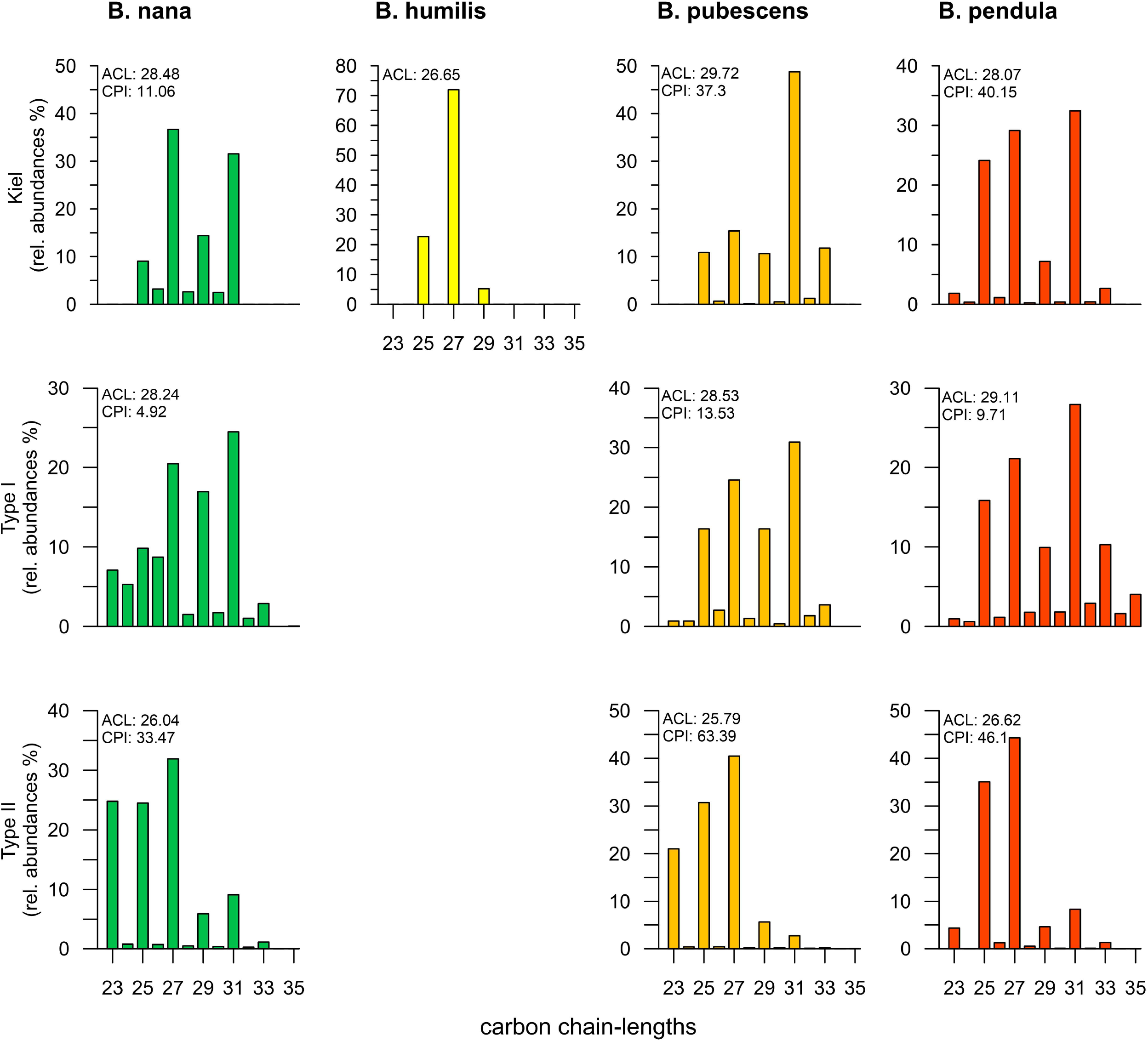
Relative abundances of *n*-alkane patterns for *Betula nana, Betula humilis, Betula pubescens* and *Betula pendula*. The samples from our study (Kiel) were compared with literature data previously divided into two subgroups (Type I and Type II; see text for details) with similar compositions. Note, to the best of our knowledge there are no previous studies reporting the epicuticular wax composition of *Betula humilis*.

**Table 2:** Relative *n*-alkane abundances of grouped *Betula* species

|  |  | Relative abundance<br>(%) |  |  |  |  |  |  |  |  |  |  |  |  |  |
| --- | --- | --- | --- | --- | --- | --- | --- | --- | --- | --- | --- | --- | --- | --- | --- |
| Species | Typ | 23 | 24 | 25 | 26 | 27 | 28 | 29 | 30 | 31 | 32 | 33 | 34 | 35 | ACL |
| *B. nana* | Kiel | 0 | 0 | 9 | 3 | 37 | 3 | 14 | 2 | 32 | 0 | 0 | 0 | 0 | 28.49 |
| *B. nana* | I | 7 | 5 | 10 | 9 | 20 | 1 | 17 | 2 | 24 | 1 | 3 | 0 | 0 | 28.24 |
| *B. nana* | II | 25 | 1 | 25 | 1 | 32 | 1 | 6 | 0 | 9 | 0 | 1 | 0 | 0 | 26.04 |
| *B. humilis* | Kiel | 0 | 0 | 23 | 0 | 72 | 0 | 5 | 0 | 0 | 0 | 0 | 0 | 0 | 26.65 |
| *B. pubescens* | Kiel | 0 | 0 | 11 | 1 | 15 | 0 | 11 | 1 | 49 | 1 | 12 | 0 | 0 | 29.72 |
| *B. pubescens* | I | 1 | 1 | 16 | 3 | 25 | 1 | 16 | 0 | 31 | 2 | 4 | 0 | 0 | 28.53 |
| *B. pubescens* | II | 21 | 0 | 31 | 0 | 40 | 0 | 6 | 0 | 3 | 0 | 0 | 0 | 0 | 25.79 |
| *B. pendula* | Kiel | 2 | 0 | 24 | 1 | 29 | 0 | 7 | 0 | 32 | 0 | 3 | 0 | 0 | 28.07 |
| *B. pendula* | I | 1 | 1 | 16 | 1 | 21 | 2 | 10 | 2 | 28 | 3 | 10 | 2 | 4 | 29.11 |
| *B. pendula* | II | 4 | 0 | 35 | 1 | 44 | 1 | 5 | 0 | 8 | 0 | 1 | 0 | 0 | 26.62 |

All *Betula* tree species from Kiel and from globally distributed type I were distinct from grasses and shrubs by a prominent prevalence of the *n*C_27_ alkane (Fig. 1 and 4). Grasses are mainly dominated by very long-chain n-alkanes with *n*C_31_ and *n*C_33_ or even *n*C_35_, and therefore are characterized by a high ACL (>30) [44,49,67]. Typical pioneer shrubs of the late glacial period such as A*rtemisia sp*. or *Junipers sp*. are also characterized by a dominance at *n*C_31_ to *n*C_35_, which is not prevalent in *Betula* species [45,91,92]. Other prominent species such as *Pinus sp*., which are probably the most widely spread conifer species in Europe, synthesize *n*-alkanes in the range from *n*C_27_ and *n*C_31_ [45,93]. However, the quantitative amounts are significantly lower, pointing to subordinate proportions of sedimentary *n*-alkanes originating from pines [47].

The type II birch species had *n*-alkane distributions similar to aquatic macrophytes and non-emergent (submerged and floating) aquatic plants, with a prevalence of mid-chain n-alkanes (*n*C_23_, *n*C_25_). These homologues can be a major source to geological archives, especially in lake sediments, often expressed in the P_aq_ proxy [54,94]. It has been postulated that terrestrial plants correspond to P_aq_ < 0.1, emergent macrophytes to P_aq_ 0.1 – 0.4 and non-emergent macrophytes to P_aq_ 0.4 – 1.0 to [54]. Each type II birch species, as well as *B. humilis* from our study, had a P_aq_ value above 0.75 corresponding to a non-emergent macrophyte. Therefore, when applying the P_aq_ proxy in the study of lake sediments receiving birch input, it must be considered that mid-chain *n*-alkanes may derive from either aquatic macrophytes or alternatively from birch trees.

*n*-Alkane abundance or chain-length based indicators like ACL have been used as a proxy for temperature, aridity, geographic location, or vapour pressure deficit [95–98]. The variation of the ACL compiled for *B. nana* did not show a trend with climatic drivers. The *B. nana* trees listed, mostly derived from cold environments such as Alaska, Greenland or Siberia and varied in their ACL between 26.4 and 28.9 [80,82,83]. Even two samples both originating from Norway varied by over 2 units in their ACL [81,84]. The *B. nana* investigated from Kiel, and therefore from the warmest region, did not reveal the highest ACL, but rather values between those of leaves from Greenland and Alaska. The ACL distributions of *B. pubescens* revealed higher values in samples from a moderate climate (Kiel, Scotland), whereas samples from colder regions (Siberia, Norway) had a lower ACL. Under certain assumptions, this can be attributed a geographical or temperature effect. The origin for the *B. pubescens* leaves from the study by Pagani et al. (2006) were not reported. Similar to *B. nana*, no latitudinal or temperature trend in the *n*-alkane distribution of *B. pendula* wax was observed.

Overall, a high variability of wax *n*-alkanes within the individual species was noted without a temperature or geographical trend. This may suggest that genetic differences between the populations control wax lipid composition, preferentially. It is conceivable that not only “pure-bred” birches of the respective species were examined in these studies, but also subspecies or varieties. For example, *Betula pubescens* has several varieties that occur naturally in a narrow space like *var. pubescens, var. fragrans, var. litwinowii* and *var. pumila* [15]. Wild hybridization may also affect leaf wax composition, whereby hybridization readily occurs between species with the same chromosome number like *B. pendula* × *B. nana* (diploid × diploid), but there are also reports of interploidy-level hybrids like *pendula* × *B. pubescens* (diploid × tetraploid) [99,100]. Therefore, future studies may address the association between ploidy level and wax lipid composition of *Betula* species to investigate species determination.

### Analytical and extraction protocols used in previous *Betula* studies

Different analytic protocols have been used to extract the plant lipids as briefly descripted above. Previous studies have shown that the length of the extraction time, as well as the solvent used, had an influence on extraction yield or lipid extract composition [37,46]. Thus, *n*-alkanes and *n*-alcohols were extracted earlier than *n*-alkanoic acids and long-chain homologues earlier than the shorter ones [101]. Jetter et al. (2008) indicated extraction yields of *n*-alkanes depended on polarity of binary solvent mixtures. Moreover, saponification upon extraction [81], hydrolyzed wax esters leading to enhanced release of bound *n*-alcohols and *n*-alkanoic acids adding to the proportion of the free homologues. Since this comparative investigation used *n*-alkane distributions from different studies with different extraction methods, the results are not unequivocally comparable. For a better comparability of future work, the influence of the extraction method on the other lipid classes including *n*-alcohols, *n*-alkanoic acids and *n*-alkyl esters should be investigated and a standard extraction protocol established.

## Conclusion

The leaves of four *Betula* species, *B. nana, B. humilis, B. pubescens, B. pendula*, which are endemic in Europe were studied, aiming to investigate their epicuticular wax lipid composition. The following conclusions can be drawn from this study. The *n*-alkane compositions in leaves of *Betula* species from Kiel were found to be specific, allowing unambiguous differentiation. *Betula* wax *n*-alcohol and *n*-alkyl ester composition allowed a distinction to be made between *B. nana, B. humilis* and the two birch trees, however the latter two cannot be easily distinguished from each other due to a similar fingerprint. The *n*-alkanoic acids seemed to be less suitable for species differentiation since all four species were dominated by the C_28_ alkanoic acid, however with variations in concentration of about two orders of magnitude. A flowchart (Fig. 5) provides a simple means for discrimination of epicuticular waxes from the four birches from Kiel University.

**Figure 5:**
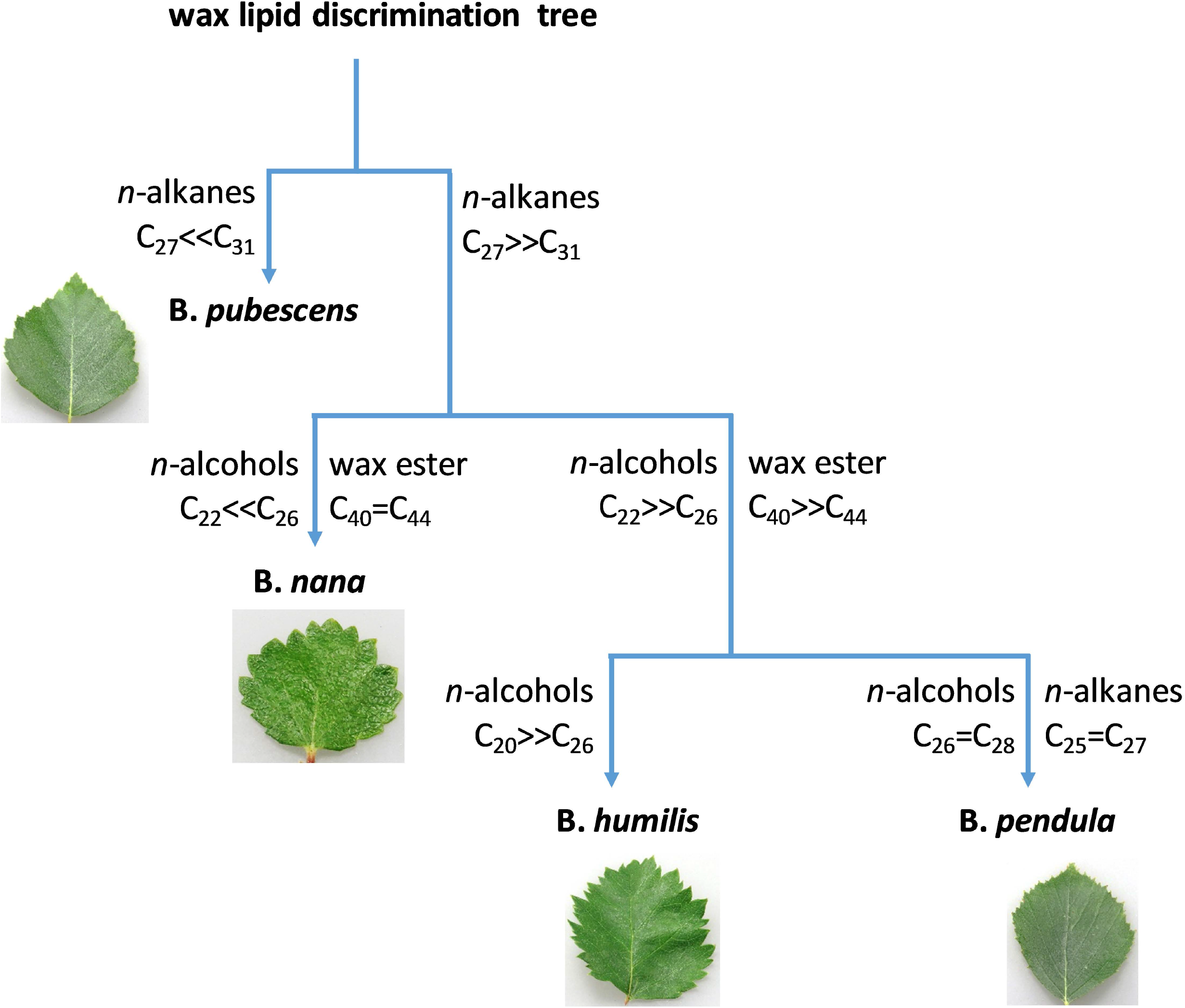
Decision making tree for differentiation of four European *Betula* species based on epicuticular wax composition

The *n*-alkyl esters consisted of different isomers with varying *n*-alcohol and *n*-alkanoic acid moieties. In the species *B. humilis* and B. *pubescens*, the dominant esterified alcohol also was the dominant free alcohol, therefore the *n*-alcohol patterns in sediments would not be disturbed by hydrolysis of the wax esters. In *B. nana* and *B. pendula* the *n*-alcohol distribution changed substantially upon ester hydrolysis, when bound homologues were released. Due to the preponderance of short-chain ester-bound alkanoic acids in wax esters, the distribution of free long-chain alkanoic acids was only slightly impaired. The ratio was influenced in *B. nana* only, as large amounts of bound C_20_ were released.

When comparing the *n*-alkane composition of the *Betula* waxes collected in Kiel with published data, no trend in geographical location or temperature could be identified. It appears that *Betula* wax composition is genetically controlled, and differences occur due to presence of plant hybrids or variants.

## Acknowledgements

T. Martens and V. Grote are thanked for laboratory assistance during lipid extraction. S. Petersen is thanked for her advice during leaf sampling in the Botanical Garden of Kiel University.

## Supporting information

**S1 Figure.**
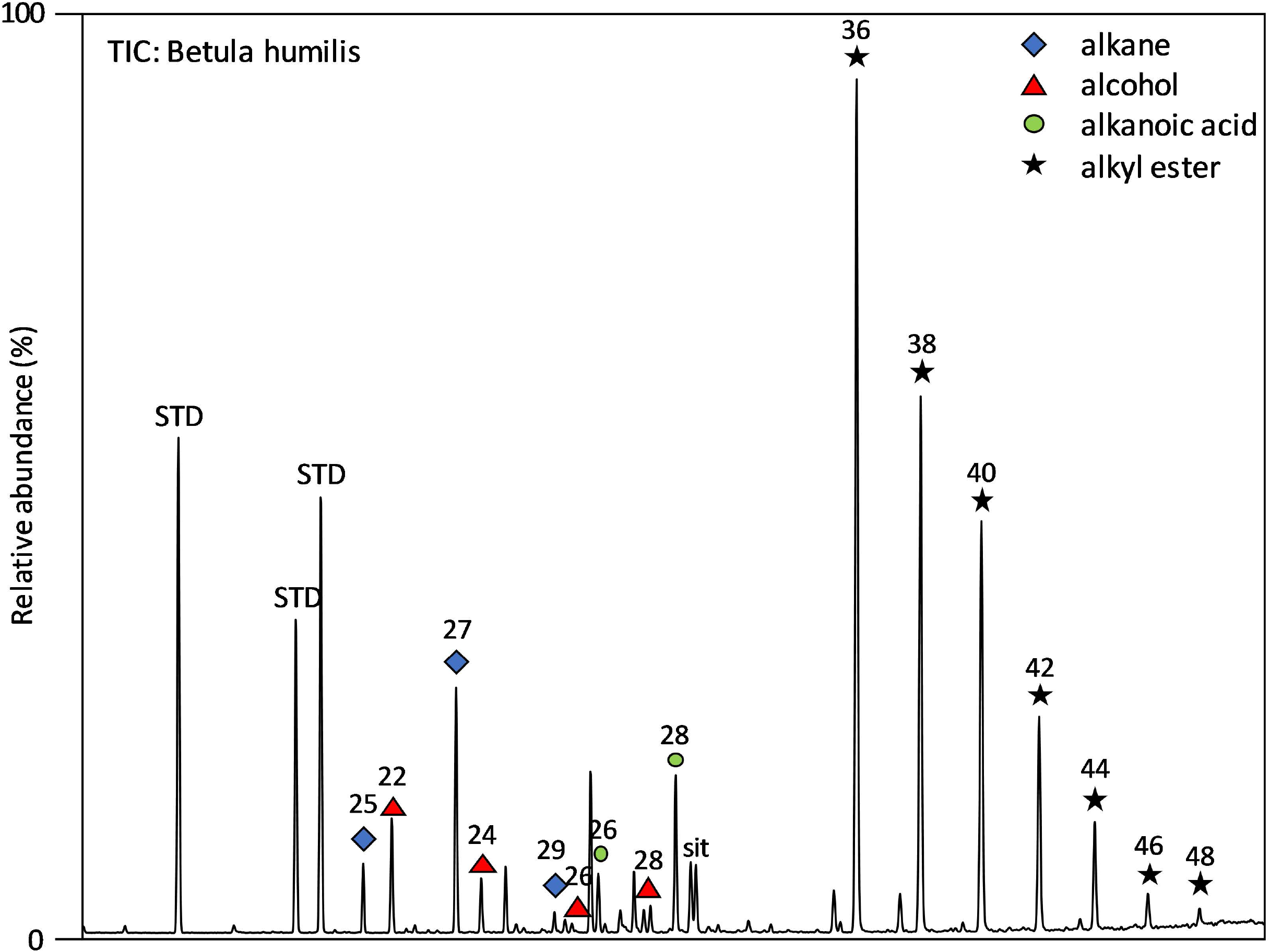
Total ion chromatogram of a *Betula humilis* leaf with the most abundant components.

**S2 Figure.**
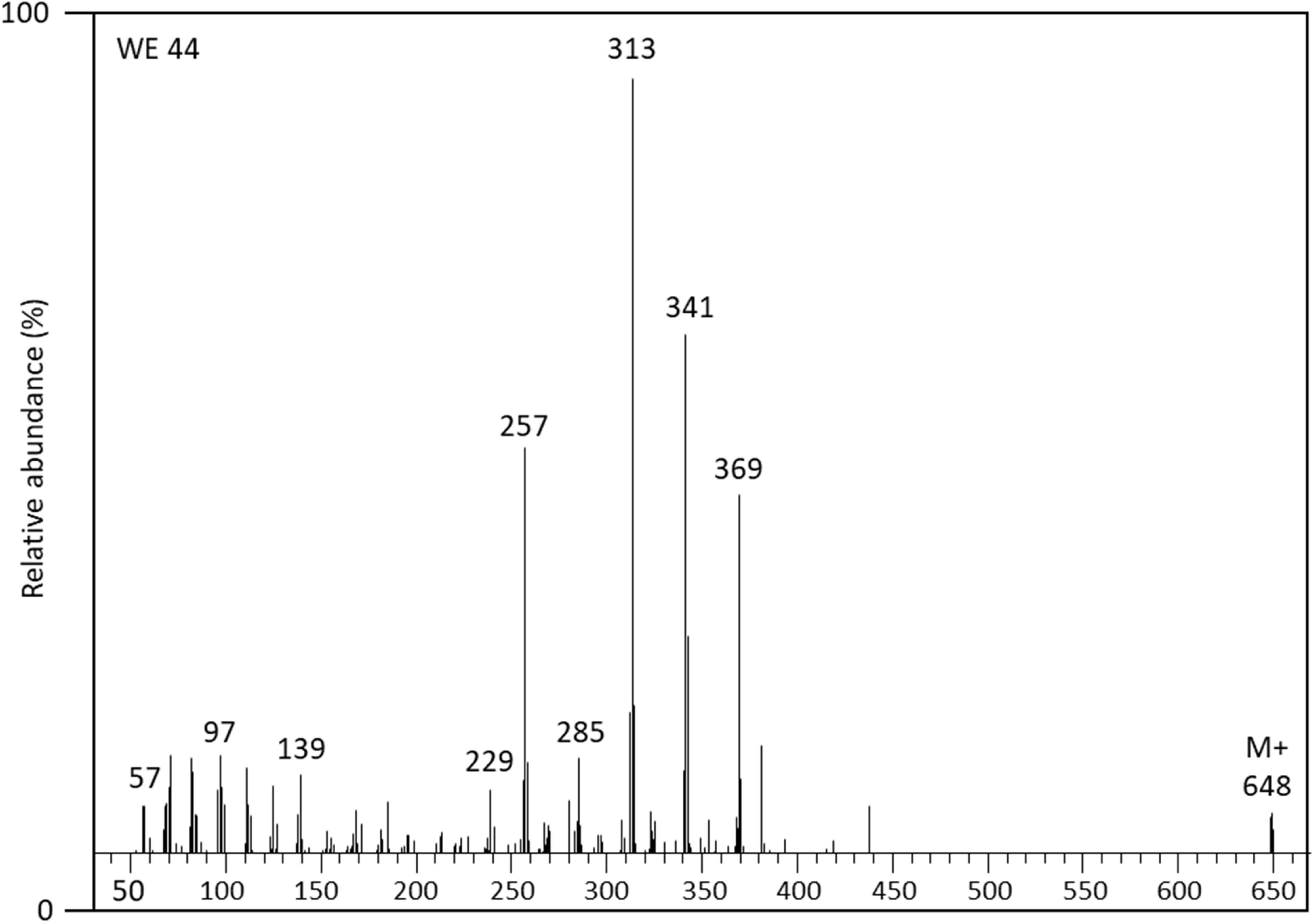
Mass spectrum of a C_44_ alkyl ester mixture (RCOOR’) of *Betula humilis*. The diagnostic ions are shown for the acid fragments (RCO_2_H_2_^+^) and the molecular ion (M^+^).

**S3 Table.**
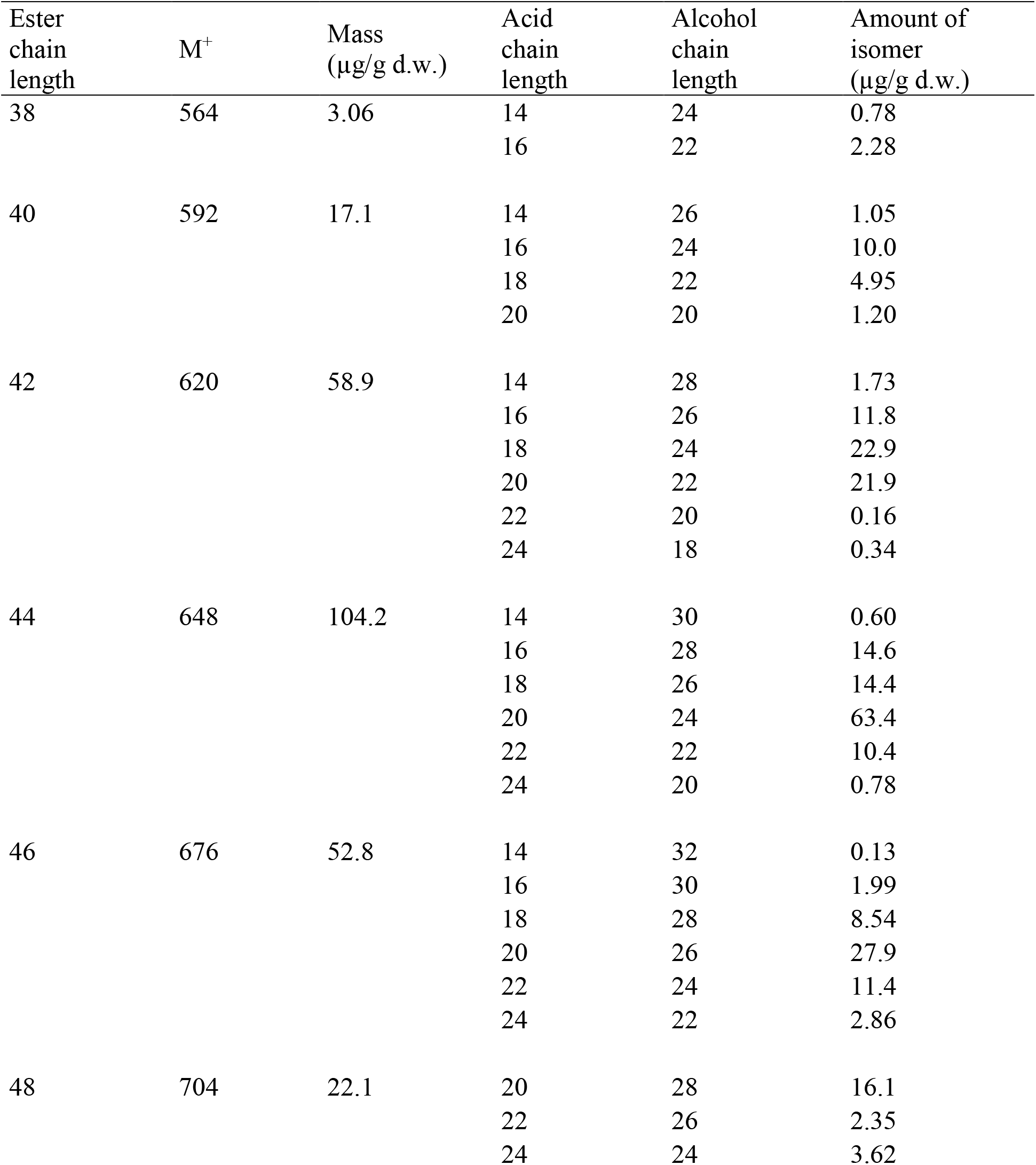

| Ester chain length | M<sup>+</sup> | Mass (μg/g d.w.) | Acid chain length | Alcohol chain length | Amount of isomer (μg/g d.w.) |
| --- | --- | --- | --- | --- | --- |
| 38 | 564 | 3.06 | 14 | 24 | 0.78 |
|  |  |  | 16 | 22 | 2.28 |
| 40 | 592 | 17.1 | 14 | 26 | 1.05 |
|  |  |  | 16 | 24 | 10.0 |
|  |  |  | 18 | 22 | 4.95 |
|  |  |  | 20 | 20 | 1.20 |
| 42 | 620 | 58.9 | 14 | 28 | 1.73 |
|  |  |  | 16 | 26 | 11.8 |
|  |  |  | 18 | 24 | 22.9 |
|  |  |  | 20 | 22 | 21.9 |
|  |  |  | 22 | 20 | 0.16 |
|  |  |  | 24 | 18 | 0.34 |
| 44 | 648 | 104.2 | 14 | 30 | 0.60 |
|  |  |  | 16 | 28 | 14.6 |
|  |  |  | 18 | 26 | 14.4 |
|  |  |  | 20 | 24 | 63.4 |
|  |  |  | 22 | 22 | 10.4 |
|  |  |  | 24 | 20 | 0.78 |
| 46 | 676 | 52.8 | 14 | 32 | 0.13 |
|  |  |  | 16 | 30 | 1.99 |
|  |  |  | 18 | 28 | 8.54 |
|  |  |  | 20 | 26 | 27.9 |
|  |  |  | 22 | 24 | 11.4 |
|  |  |  | 24 | 22 | 2.86 |
| 48 | 704 | 22.1 | 20 | 28 | 16.1 |
|  |  |  | 22 | 26 | 2.35 |
|  |  |  | 24 | 24 | 3.62 |

**S4 Table.**

| Ester chain length | M+ | Mass (µg/g d.w.) | Acid chain length | Alcohol chain length | Amount of isomer (µg/g d.w.) |
| --- | --- | --- | --- | --- | --- |
| 36 | 536 | 1047 | 14 | 22 | 6.5 |
|  |  |  | 16 | 20 | 1040 |
| 38 | 564 | 601 | 14 | 24 | 7.5 |
|  |  |  | 16 | 22 | 420 |
|  |  |  | 18 | 20 | 173 |
| 40 | 592 | 532 | 14 | 26 | 3.0 |
|  |  |  | 16 | 24 | 254 |
|  |  |  | 18 | 22 | 123 |
|  |  |  | 20 | 20 | 150 |
| 42 | 620 | 310 | 14 | 28 | 2.4 |
|  |  |  | 16 | 26 | 57.7 |
|  |  |  | 18 | 24 | 57.9 |
|  |  |  | 20 | 22 | 147 |
|  |  |  | 22 | 20 | 44.7 |
| 44 | 648 | 180 | 16 | 28 | 39.7 |
|  |  |  | 18 | 26 | 10.9 |
|  |  |  | 20 | 24 | 59.2 |
|  |  |  | 22 | 22 | 49.3 |
|  |  |  | 24 | 20 | 21.1 |
| 46 | 676 | 70 | 16 | 30 | 6.9 |
|  |  |  | 18 | 28 | 5.6 |
|  |  |  | 20 | 26 | 10.2 |
|  |  |  | 22 | 24 | 16.6 |
|  |  |  | 24 | 22 | 11.3 |
|  |  |  | 26 | 20 | 19.8 |
| 48 | 704 | 35 | 16 | 32 | 3.62 |
|  |  |  | 20 | 28 | 6.03 |
|  |  |  | 24 | 24 | 4.95 |
|  |  |  | 26 | 22 | 6.18 |
|  |  |  | 28 | 20 | 14.2 |

**S5 Table.**

| Ester chain length | M+ | Mass (µg/g d.w.) | Acid chain length | Alcohol chain length | Amount of isomer (µg/g d.w.) |
| --- | --- | --- | --- | --- | --- |
| 38 | 564 | 535.74 | 14 | 24 | 13.6 |
|  |  |  | 16 | 22 | 519 |
|  |  |  | 18 | 20 | 2.73 |
|  |  |  | 20 | 18 | 0.57 |
| 40 | 592 | 290.49 | 14 | 26 | 7.29 |
|  |  |  | 16 | 24 | 208 |
|  |  |  | 18 | 22 | 71.5 |
|  |  |  | 20 | 20 | 4.06 |
| 42 | 620 | 147.36 | 14 | 28 | 4.85 |
|  |  |  | 16 | 26 | 67.3 |
|  |  |  | 18 | 24 | 22.8 |
|  |  |  | 20 | 22 | 52.4 |
| 44 | 648 | 68.44 | 16 | 28 | 39.7 |
|  |  |  | 18 | 26 | 9.65 |
|  |  |  | 20 | 24 | 19.0 |

**S6 Table.**
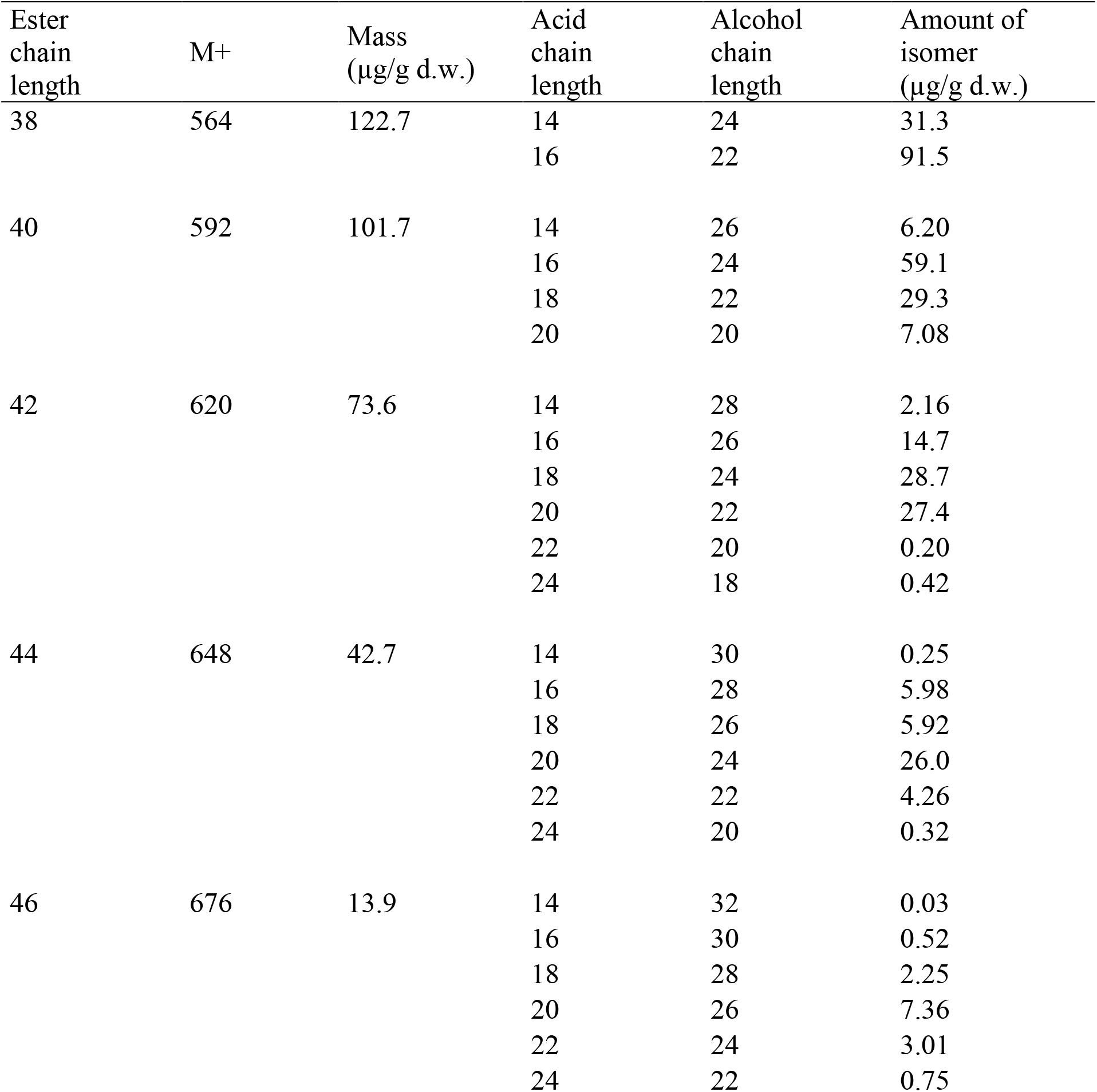

| Ester chain length | M+ | Mass (µg/g d.w.) | Acid chain length | Alcohol chain length | Amount of isomer (µg/g d.w.) |
| --- | --- | --- | --- | --- | --- |
| 38 | 564 | 122.7 | 14 | 24 | 31.3 |
|  |  |  | 16 | 22 | 91.5 |
| 40 | 592 | 101.7 | 14 | 26 | 6.20 |
|  |  |  | 16 | 24 | 59.1 |
|  |  |  | 18 | 22 | 29.3 |
|  |  |  | 20 | 20 | 7.08 |
| 42 | 620 | 73.6 | 14 | 28 | 2.16 |
|  |  |  | 16 | 26 | 14.7 |
|  |  |  | 18 | 24 | 28.7 |
|  |  |  | 20 | 22 | 27.4 |
|  |  |  | 22 | 20 | 0.20 |
|  |  |  | 24 | 18 | 0.42 |
| 44 | 648 | 42.7 | 14 | 30 | 0.25 |
|  |  |  | 16 | 28 | 5.98 |
|  |  |  | 18 | 26 | 5.92 |
|  |  |  | 20 | 24 | 26.0 |
|  |  |  | 22 | 22 | 4.26 |
|  |  |  | 24 | 20 | 0.32 |
| 46 | 676 | 13.9 | 14 | 32 | 0.03 |
|  |  |  | 16 | 30 | 0.52 |
|  |  |  | 18 | 28 | 2.25 |
|  |  |  | 20 | 26 | 7.36 |
|  |  |  | 22 | 24 | 3.01 |
|  |  |  | 24 | 22 | 0.75 |

## References

1. Crane PR, Stockey RA. Betula leaves and reproductive structures from the Middle Eocene of British Columbia, Canada. Can J Bot. 1987;65: 2490–2500. doi:10.1139/b87-338

2. Järvinen P, Palme A, Orlando Morales L, Lannenpaa M, Keinanen M, Sopanen T, et al. Phylogenetic relationships of Betula species (Betulaceae) based on nuclear ADH and chloroplast matK sequences. Am J Bot. 2004;91: 1834–1845. doi:10.3732/ajb.91.11.1834

3. Schenk MF, Thienpont C-N, Koopman WJM, Gilissen LJWJ, Smulders MJM. Phylogenetic relationships in Betula (Betulaceae) based on AFLP markers. Tree Genet Genomes. 2008;4: 911. doi: 10.1007/s11295-008-0162-0

4. Bina H, Yousefzadeh H, Ali SS, Esmailpour M. Phylogenetic relationships, molecular taxonomy, biogeography of Betula, with emphasis on phylogenetic position of Iranian populations. Tree Genet Genomes. 2016; 12: 84. doi:10.1007/s11295-016-1037-4

5. Julkunen-Tiitto R, Rousi M, Bryant J, Sorsa S, Keinänen M, Sikanen H. Chemical diversity of several Betulaceae species: comparison of phenolics and terpenoids in northern birch stems. Trees. 1996;11: 16. doi:10.1007/s004680050053

6. Keinänen M, Julkunen-Tiitto R, Rousi M, Tahvanainen J. Taxonomic implications of phenolic variation in leaves of birch (Betula L.) species. Biochem Syst Ecol. 1999;27: 243–254. doi:10.1016/S0305-1978(98)00086-6

7. Orav A, Arak E, Boikova T, Raal A. Essential oil in Betula spp. leaves naturally growing in Estonia. Biochem Syst Ecol. 2011;39: 744–748. doi:10.1016/j.bse.2011.06.013

8. Depciuch J, Kasprzyk I, Drzymała E, Parlinska-Wojtan M. Identification of birch pollen species using FTIR spectroscopy. Aerobiologia (Bologna). 2018;34: 525–538. doi:10.1007/s10453-018-9528-4

9. Furlow JJ. The genera of Betulaceae in the southeastern United States. J Arnold Arboretum. 1990;71: 1–67. doi:10.5962/bhl.part.24925

10. Maliouchenko O, Palmé AE, Buonamici A, Vendramin GG, Lascoux M. Comparative phylogeography and population structure of European Betula species, with particular focus on B. pendula and B. pubescens. J Biogeogr. 2007;34: 1601–1610. doi: 10.1111/j.1365-2699.2007.01729.x

11. Wang N, McAllister HA, Bartlett PR, Buggs RJA. Molecular phylogeny and genome size evolution of the genus Betula (Betulaceae). Ann Bot. 2016;117: 1023–1035. doi:10.1093/aob/mcw048

12. Winkler H. Betulaceae Das Pflanzenreich. 39th ed. Engler A, editor. 1904.

13. De Jong PC. An introduction to Betula: Its morphology, evolution, classification and distribution with a survey of recent work. IDS Betula Symp Int Dendrol Soc Susses, UK. 1993.

14. Skvortsov AK. A new system of the genus Betula L. Bull Mosc Natur Soc. 2002;107: 73–76.

15. Ashburner K, McAllister HA. The Genus Betula: A Taxonomic Revision of Birches. 2013.

16. Freund H, Birks HH, Birks HJB. The identification of wingless Betula fruits in Weichselian sediments in the Gross Todtshorn borehole (Lower Saxony, Germany) – the occurrence of Betula humilis Schrank. Veg Hist Archaeobot. 2001;10: 107–115. doi:10.1007/PL00006919

17. van Dinter M, Birks HH. Distinguishing fossil Betula nana and B. pubescens using their wingless fruits: implications for the late-glacial vegetational history of western Norway. Veg Hist Archaeobot. 1996;5: 229–240. doi:10.1007/BF00217500

18. Tsuda Y, Semerikov V, Sebastiani F, Vendramin GG, Lascoux M. Multispecies genetic structure and hybridization in the Betula genus across Eurasia. Mol Ecol. 2017;26: 589–605. doi:10.1111/mec.13885

19. Thomson AM, Dick CW, Pascoini AL, Dayanandan S. Despite introgressive hybridization, North American birches (Betula spp.) maintain strong differentiation at nuclear microsatellite loci. Tree Genet Genomes. 2015;11: 101. doi:10.1007/s11295-015-0922-6

20. Thórsson AT, Salmela E, Anamthawat-Jónsson K. Morphological, cytogenetic, and molecular evidence for introgressive hybridization in birch. J Hered. 2001;92: 404–8. doi:10.1093/jhered/92.5.404

21. Tralau H. The recent and fossil distribution of some boreal and arctic montane plants in Europe. Ark. Bot. Ser. 2; 1963.

22. Krüger S, Damrath M. In search of the Bølling-Oscillation: a new high resolution pollen record from the locus classicus Lake Bølling, Denmark. Veg Hist Archaeobot. 2019; doi:10.1007/s00334-019-00736-3

23. Eusterhues K, Lechterbeck J, Schneider J, Wolf-Brozio U. Late- and Post-Glacial evolution of Lake Steisslingen. Palaeogeogr Palaeoclimatol Palaeoecol. 2002;187: 341–371. doi:10.1016/S0031-0182(02)00486-8

24. Lotter AF. Late-glacial and Holocene vegetation history and dynamics as shown by pollen and plant macrofossil analyses in annually laminated sediments from Soppensee, central Switzerland. Veg Hist Archaeobot. 1999;8: 165–184. doi:10.1007/BF02342718

25. Lotter AF, Eicher U, Siegenthaler U, Birks HJB. Late-glacial climatic oscillations as recorded in Swiss lake sediments. J Quat Sci. 1992;7: 187–204. doi:10.1002/jqs.3390070302

26. Veski S, Amon L, Heinsalu A, Reitalu T, Saarse L, Stivrins N, et al. Lateglacial vegetation dynamics in the eastern Baltic region between 14,500 and 11,400calyrBP: A complete record since the Bølling (GI-1e) to the Holocene. Quat Sci Rev. 2012;40: 39–53. doi:10.1016/j.quascirev.2012.02.013

27. Tarasov PE, Bezrukova E V., Krivonogov SK. Late Glacial and Holocene changes in vegetation cover and climate in southern Siberia derived from a 15 kyr long pollen record from Lake Kotokel. Clim Past. 2009;5: 285–295. doi:10.5194/cp-5-285-2009

28. Mortensen MF, Birks HH, Christensen C, Holm J, Noe-Nygaard N, Odgaard BV, et al. Lateglacial vegetation development in Denmark – New evidence based on macrofossils and pollen from Slotseng, a small-scale site in southern Jutland. Quat Sci Rev. 2011;30: 2534–2550. doi:10.1016/j.quascirev.2011.04.018

29. Rösch M, Lechterbeck J. Seven Millennia of human impact as reflected in a high resolution pollen profile from the profundal sediments of Litzelsee, Lake Constance region, Germany. Veg Hist Archaeobot. 2016;25: 339–358. doi:10.1007/s00334-015-0552-9

30. Jahns S. Late-glacial and Holocene woodland dynamics and land-use history of the Lower Oder valley, north-eastern Germany, based on two, AMS14C-dated, pollen profiles. Veg Hist Archaeobot. 2000;9: 111–123. doi:10.1007/BF01300061

31. Birks HJB. The Identification of Betula nana Pollen. New Phytol. 1968;67: 309–314. doi:10.1111/j.1469-8137.1968.tb06386.x

32. Mäkelä EM. Size distinctions between Betula pollen types — A review. Grana. 1996;35: 248–256. doi: 10.1080/00173139609430011

33. Caseldine C. Changes in Betula in the Holocene record from Iceland—a palaeoclimatic record or evidence for early Holocene hybridisation? Rev Palaeobot Palynol. 2001;117: 139–152. doi:10.1016/S0034-6667(01)00082-3

34. Karlsdóttir L, Thórsson ÆT, Hallsdóttir M, Sigurgeirsson A, Eysteinsson T, Anamthawat-Jónsson K. Differentiating pollen of Betula species from Iceland. Grana. 2007;46: 78–84. doi:10.1080/00173130701237832

35. de Klerk P, Theuerkauf M, Joosten H. Vegetation, recent pollen deposition, and distribution of some non-pollen palynomorphs in a degrading ice-wedge polygon mire complex near Pokhodsk (NE Siberia), including size-frequency analyses of pollen attributable to Betula. Rev Palaeobot Palynol. 2017;238: 122–143. doi:10.1016/j.revpalbo.2016.11.015

36. Jenks MA, Joly RJ, Peters PJ, Rich PJ, Axtell JD, Ashworth EN. Chemically Induced Cuticle Mutation Affecting Epidermal Conductance to Water Vapor and Disease Susceptibility in Sorghum bicolor (L.) Moench. Plant Physiol. 1994; 105: 1239–1245. doi:10.1104/pp.105.4.1239

37. Jetter R, Kunst L. Plant surface lipid biosynthetic pathways and their utility for metabolic engineering of waxes and hydrocarbon biofuels. Plant J. 2008;54: 670–683. doi: 10.1111/j.1365-313X.2008.03467.x

38. Sieber P, Schorderet M, Ryser U, Buchala A, Kolattukudy P, Métraux J-P, et al. Transgenic Arabidopsis Plants Expressing a Fungal Cutinase Show Alterations in the Structure and Properties of the Cuticle and Postgenital Organ Fusions. Plant Cell. 2000;12: 721–737. doi:10.1105/tpc.12.5.721

39. Long LM, Patel HP, Cory WC, Stapleton AE. The maize epicuticular wax layer provides UV protection. Funct Plant Biol. 2003;30: 75. doi:10.1071/FP02159

40. Eglinton G, Hamilton RJ. Leaf Epicuticular Waxes. Science (80-). 1967;156: 1322–1335. doi:10.1126/science.156.3780.1322

41. Eglinton G, Gonzalez AG, Hamilton RJ, Raphael RA. Hydrocarbon constituents of the wax coatings of plant leaves: A taxonomic survey. Phytochemistry. 1962;1: 89–102. doi:10.1016/S0031-9422(00)88006-1

42. Herbin GA, Robins PA. Patterns of variation and development in leaf wax alkanes. Phytochemistry. 1969;8: 1985–1998. doi:10.1016/S0031-9422(00)88085-1

43. Gülz P-G. Epicuticular Leaf Waxes in the Evolution of the Plant Kingdom. J Plant Physiol. 1994; 143: 453–464. doi: 10.1016/S0176-1617(11)81807-9

44. Maffei M. Chemotaxonomic significance of leaf wax alkanes in the gramineae. Biochem Syst Ecol. 1996;24: 53–64. doi:10.1016/0305-1978(95)00102-6

45. Schwark L, Zink K, Lechterbeck J. Reconstruction of postglacial to early Holocene vegetation history in terrestrial Central Europe via cuticular lipid biomarkers and pollen records from lake sediments. Geology. 2002;30: 463. doi:10.1130/0091-7613(2002)030<0463:ROPTEH>2.0.CO;2

46. Buschhaus C, Herz H, Jetter R. Chemical Composition of the Epicuticular and Intracuticular Wax Layers on Adaxial Sides of Rosa canina Leaves. Ann Bot. 2007;100: 1557–1564. doi:10.1093/aob/mcm255

47. Diefendorf AF, Freeman KH, Wing SL, Graham H V. Production of n-alkyl lipids in living plants and implications for the geologic past. Geochim Cosmochim Acta. 2011;75: 7472–7485. doi: 10.1016/j.gca.2011.09.028

48. Diefendorf AF, Leslie AB, Wing SL. Leaf wax composition and carbon isotopes vary among major conifer groups. Geochim Cosmochim Acta. 2015; 170: 145–156. doi:10.1016/j.gca.2015.08.018

49. Bush RT, McInerney FA. Leaf wax n-alkane distributions in and across modern plants: Implications for paleoecology and chemotaxonomy. Geochim Cosmochim Acta. 2013;117: 161–179. doi: 10.1016/j.gca.2013.04.016

50. Mueller-Niggemann C, Schwark L. Chemotaxonomy and diagenesis of aliphatic hydrocarbons in rice plants and soils from land reclamation areas in the Zhejiang Province, China. Org Geochem. 2015; 83-84: 215–226. doi:10.1016/j.orggeochem.2015.03.016

51. Guo Y, Li JJ, Busta L, Jetter R. Coverage and composition of cuticular waxes on the fronds of the temperate ferns Pteridium aquilinum, Cryptogramma crispa, Polypodium glycyrrhiza, Polystichum munitum and Gymnocarpium dryopteris. Ann Bot. 2018;122: 555–568. doi:10.1093/aob/mcy078

52. Blumer M, Guillard RRL, Chase T. Hydrocarbons of marine phytoplankton. Mar Biol. 1971;8: 183–189. doi: 10.1007/BF00355214

53. Cranwell PA, Eglinton G, Robinson N. Lipids of aquatic organisms as potential contributors to lacustrine sediments-II. Org Geochem. 1987; 11: 513–527. doi:10.1016/0146-6380(87)90007-6

54. Ficken K., Li B, Swain D., Eglinton G. An n-alkane proxy for the sedimentary input of submerged/floating freshwater aquatic macrophytes. Org Geochem. 2000;31: 745–749. doi:10.1016/S0146-6380(00)00081-4

55. Pancost RD, Baas M, van Geel B, Sinninghe Damsté JS. Biomarkers as proxies for plant inputs to peats: an example from a sub-boreal ombrotrophic bog. Org Geochem. 2002;33: 675–690. doi: 10.1016/S0146-6380(02)00048-7

56. Shepherd T, Robertson GW, Griffiths DW, Birch ANE. Epicuticular wax ester and triacylglycerol composition in relation to aphid infestation and resistance in red raspberry (Rubus idaeus L.). Phytochemistry. 1999;52: 1255–1267. doi:10.1016/S0031-9422(99)00414-8

57. Schouten S, Woltering M, Rijpstra WIC, Sluijs A, Brinkhuis H, Sinninghe Damsté JS. The Paleocene-Eocene carbon isotope excursion in higher plant organic matter: Differential fractionation of angiosperms and conifers in the Arctic. Earth Planet Sci Lett. 2007;258: 581–592. doi: 10.1016/j.epsl.2007.04.024

58. Smith F, Wing S, Freeman K. Magnitude of the carbon isotope excursion at the Paleocene–Eocene thermal maximum: The role of plant community change. Earth Planet Sci Lett. 2007;262: 50–65. doi:10.1016/j.epsl.2007.07.021

59. Schellekens J, Buurman P. Geoderma n-Alkane distributions as palaeoclimatic proxies in ombrotrophic peat: The role of decomposition and dominant vegetation. Geoderma. 2011;164: 112–121. doi:10.1016/j.geoderma.2011.05.012

60. Jansen B, Wiesenberg GLB. Opportunities and limitations related to the application of plant-derived lipid molecular proxies in soil science. SOIL. 2017;3: 211–234. doi:10.5194/soil-3-211-2017

61. Jansen B, de Boer EJ, Cleef AM, Hooghiemstra H, Moscol-Olivera M, Tonneijck FH, et al. Reconstruction of late Holocene forest dynamics in northern Ecuador from biomarkers and pollen in soil cores. Palaeogeogr Palaeoclimatol Palaeoecol. 2013;386: 607–619. doi:10.1016/j.palaeo.2013.06.027

62. Wiesenberg GLB, Andreeva DB, Chimitdorgieva GD, Erbajeva MA, Zech W. Reconstruction of environmental changes during the late glacial and Holocene reflected in a soil-sedimentary sequence from the lower Selenga River valley, Lake Baikal region, Siberia, assessed by lipid molecular proxies. Quat Int. 2015;365: 190–202. doi:10.1016/j.quaint.2015.01.042

63. Lockheart MJ, van Bergen PF, Evershed RP. Chemotaxonomic classification of fossil leaves from the Miocene Clarkia lake deposit, Idaho, USA based on n-alkyl lipid distributions and principal component analyses. Org Geochem. 2000;31: 1223–1246. doi:10.1016/S0146-6380(00)00107-8

64. Huang Y, Lockheart MJ, Collister JW, Eglinton G. Molecular and isotopic biogeochemistry of the Miocene Clarkia Formation: hydrocarbons and alcohols. Org Geochem. 1995;23: 785–801. doi:10.1016/0146-6380(95)80001-8

65. Poynter J, Eglinton G. Molecular Composition of Three Sediments from Hole 717C: The Bengal Fan. Proceedings of the Ocean Drilling Program, 116 Scientific Results. Ocean Drilling Program; 1990. pp. 155–161. doi:10.2973/odp.proc.sr.116.151.1990

66. Castañeda IS, Werne JP, Johnson TC, Filley TR. Late Quaternary vegetation history of southeast Africa: The molecular isotopic record from Lake Malawi. Palaeogeogr Palaeoclimatol Palaeoecol. 2009;275: 100–112. doi:10.1016/j.palaeo.2009.02.008

67. Rommerskirchen F, Plader A, Eglinton G, Chikaraishi Y, Rullkötter J. Chemotaxonomic significance of distribution and stable carbon isotopic composition of long-chain alkanes and alkan-1-ols in C4 grass waxes. Org Geochem. 2006;37: 1303–1332. doi:10.1016/j.orggeochem.2005.12.013

68. Otto A, Simpson MJ. Degradation and Preservation of Vascular Plant-derived Biomarkers in Grassland and Forest Soils from Western Canada. Biogeochemistry. 2005;74: 377–409. doi:10.1007/s10533-004-5834-8

69. Han J, Calvin M. Hydrocarbon Distribution of Algae and Bacteria, and Microbiological Activity in Sediments. Proc Natl Acad Sci. 1969;64: 436–443. doi:10.1073/pnas.64.2.436

70. Jansen B, Nierop KGJ, Hageman JA, Cleef AM, Verstraten JM. The straight-chain lipid biomarker composition of plant species responsible for the dominant biomass production along two altitudinal transects in the Ecuadorian Andes. Org Geochem. 2006;37: 1514–1536. doi:10.1016/j.orggeochem.2006.06.018

71. Schäfer IK, Lanny V, Franke J, Eglinton TI, Zech M, Vysloužilová B, et al. Leaf waxes in litter and topsoils along a European transect. SOIL. 2016;2: 551–564. doi:10.5194/soil-2-551-2016

72. Franich RA, Goodin SJ, Volkman JK. Alkyl esters from pinus radiata foliage epicuticular wax. Phytochemistry. 1985;24: 2949–2952. doi:10.1016/0031-9422(85)80033-9

73. Sümmchen P, Markstädter C, Wienhaus O. Composition of the Epicuticular Wax Esters of Picea abies (L.) Karst. Zeitschrift für Naturforsch C. 1995;50: 11–14. doi:10.1515/znc-1995-1-203

74. Koch K, Ensikat H-J. The hydrophobic coatings of plant surfaces: Epicuticular wax crystals and their morphologies, crystallinity and molecular self-assembly. Micron. 2008;39: 759–772. doi:10.1016/j.micron.2007.11.010

75. Cranwell PA, Volkman JK. Alkyl and steryl esters in a recent lacustrine sediment. Chem Geol. 1981;32: 29–43. doi:10.1016/0009-2541(81)90126-1

76. van Bergen PF, Bull ID, Poulton PR, Evershed RP. Organic geochemical studies of soils from the Rothamsted Classical Experiments—I. Total lipid extracts, solvent insoluble residues and humic acids from Broadbalk Wilderness. Org Geochem. 1997;26: 117–135. doi:10.1016/S0146-6380(96)00134-9

77. Lihavainen J, Ahonen V, Keski-Saari S, Sõber A, Oksanen E, Keinänen M. Low vapor pressure deficit reduces glandular trichome density and modifies the chemical composition of cuticular waxes in silver birch leaves. Tree Physiol. 2017;37: 1166–1181. doi:10.1093/treephys/tpx045

78. Zech M, Andreev A, Zech R, Müller S, Hambach U, Frechen M, et al. Quaternary vegetation changes derived from a loess-like permafrost palaeosol sequence in northeast Siberia using alkane biomarker and pollen analyses. Boreas. 2010;39: 540–550. doi: 10.1111/j.1502-3885.2009.00132.x

79. Tarasov PE, Müller S, Zech M, Andreeva D, Diekmann B, Leipe C. Last glacial vegetation reconstructions in the extreme-continental eastern Asia: Potentials of pollen and n-alkane biomarker analyses. Quat Int. 2013;290–291: 253–263. doi:10.1016/j.quaint.2012.04.007

80. Berke MA, Cartagena Sierra A, Bush R, Cheah D, O’Connor K. Controls on leaf wax fractionation and δ2H values in tundra vascular plants from western Greenland. Geochim Cosmochim Acta. 2019;244: 565–583. doi:10.1016/j.gca.2018.10.020

81. Mayes RW, Beresford NA, Lamb CS, Barnett CL, Howard BJ, Jones B-EV, et al. Novel approaches to the estimation of intake and bioavailability of radiocaesium in ruminants grazing forested areas. Sci Total Environ. 1994; 157: 289–300. doi:10.1016/0048-9697(94)90592-4

82. Daniels WC, Russell JM, Giblin AE, Welker JM, Klein ES, Huang Y. Hydrogen isotope fractionation in leaf waxes in the Alaskan Arctic tundra. Geochim Cosmochim Acta. 2017;213: 216–236. doi:10.1016/j.gca.2017.06.028

83. Ronkainen T, Väliranta M, McClymont E, Biasi C, Salonen S, Fontana S, et al. A combined biogeochemical and palaeobotanical approach to study permafrost environments and past dynamics. J Quat Sci. 2015;30: 189–200. doi:10.1002/jqs.2763

84. Balascio NL, D’Andrea WJ, Gjerde M, Bakke J. Hydroclimate variability of High Arctic Svalbard during the Holocene inferred from hydrogen isotopes of leaf waxes. Quat Sci Rev. 2018; 183: 177–187. doi:10.1016/j.quascirev.2016.11.036

85. Rao SJ, Iason GR, Hulbert IA, Mayes RW, Racey PA. Estimating diet composition for mountain hares in newly established native woodland: development and application of plant-wax faecal markers. Can J Zool. 2003;81: 1047–1056. doi:10.1139/z03-093

86. Pagani M, Pedentchouk N, Huber M, Sluijs A, Schouten S, Brinkhuis H, et al. Arctic hydrology during global warming at the Palaeocene/Eocene thermal maximum. Nature. 2006;442: 671–675. doi:10.1038/nature05043

87. Sachse D, Radke J, Gleixner G. δD values of individual n-alkanes from terrestrial plants along a climatic gradient – Implications for the sedimentary biomarker record. Org Geochem. 2006;37: 469–483. doi:10.1016/j.orggeochem.2005.12.003

88. van den Bos V, Engels S, Bohncke SJP, Cerli C, Jansen B, Kalbitz K, et al. Late Holocene changes in vegetation and atmospheric circulation at Lake Uddelermeer (The Netherlands) reconstructed using lipid biomarkers and compound-specific δD analysis. J Quat Sci. 2018;33: 100–111. doi: 10.1002/jqs.3006

89. Dawson LA, Towers W, Mayes RW, Craig J, Väisänen RK, Waterhouse EC. The use of plant hydrocarbon signatures in characterizing soil organic matter. Geol Soc London, Spec Publ. 2004;232: 269–276. doi:10.1144/GSL.SP.2004.232.01.24

90. Huang Y, Eglinton G, Ineson P, Bol R, Harkness DD. The effects of nitrogen fertilisation and elevated CO2 on the lipid biosynthesis and carbon isotopic discrimination in birch seedlings (Betula pendula). Plant Soil. 1999;216: 35–45. doi:10.1023/A:1004771431093

91. Maffei M, Badino S, Bossi S. Chemotaxonomic significance of leaf wax n-alkanes in the Pinales (Coniferales). J Biol Res (Thessaloniki, Greece). 2004;1: 3–19. Available: http://www.auth.gr/jbr/papers20041/01-2004.pdf

92. Rajčević N, Janaćković P, Dodoš T, Tešević V, Marin PD. Biogeographic Variation of Foliar n-Alkanes of Juniperus communis var. saxatilis Pallas from the Balkans. Chem Biodivers. 2014; 11: 1923–1938. doi:10.1002/cbdv.201400048

93. Ali HAM, Mayes RW, Hector BL, Verma AK, Ørskov ER. The possible use of n-alkanes, long-chain fatty alcohols and long-chain fatty acids as markers in studies of the botanical composition of the diet of free-ranging herbivores. J Agric Sci. 2005;143: 85–95. doi:10.1017/S0021859605004958

94. Aichner B, Herzschuh U, Wilkes H. Influence of aquatic macrophytes on the stable carbon isotopic signatures of sedimentary organic matter in lakes on the Tibetan Plateau. Org Geochem. 2010; 41: 706–718. doi:10.1016/j.orggeochem.2010.02.002

95. Hoffmann B, Kahmen A, Cernusak LA, Arndt SK, Sachse D. Abundance and distribution of leaf wax n-alkanes in leaves of Acacia and Eucalyptus trees along a strong humidity gradient in northern Australia. Org Geochem. 2013;62: 62–67. doi:10.1016/j.orggeochem.2013.07.003

96. Eley YL, Hren MT. Reconstructing vapor pressure deficit from leaf wax lipid molecular distributions. Sci Rep. 2018;8: 3967. doi:10.1038/s41598-018-21959-w

97. Tipple BJ, Pagani M. Environmental control on eastern broadleaf forest species’ leaf wax distributions and d/h ratios. Geochim Cosmochim Acta. 2013;111: 64–77. doi:10.1016/j.gca.2012.10.042

98. Kirkels FMSA, Jansen B, Kalbitz K. Consistency of plant-specific n-alkane patterns in plaggen ecosystems: A review. The Holocene. 2013;23: 1355–1368. doi:10.1177/0959683613486943

99. Elkington TT. Introgressive hybridization between Betula nana L. and B. pubescens Ehrh. in North-West Iceland. New Phytol. 1968;67: 109–118. doi:10.1111/j.1469-8137.1968.tb05459.x

100. Palme AE, Su Q, Palsson S, Lascoux M. Extensive sharing of chloroplast haplotypes among European birches indicates hybridization among Betula pendula, B. pubescens and B. nana. Mol Ecol. 2004;13: 167–178. doi:10.1046/j.1365-294X.2003.02034.x

101. Stammitti L, Derridj S, Garrec JP. Leaf epicuticular lipids of Prunus laurocerasus: importance of extraction methods. Phytochemistry. 1996;43: 45–48. doi:10.1016/0031-9422(96)00241-5

